# A Deep Feature Learning Approach for Mapping the Brain’s Microarchitecture and Organization

**DOI:** 10.1101/2020.05.26.117473

**Authors:** Aishwarya H. Balwani, Eva L. Dyer

**Affiliations:** AHB is in the School of Electrical & Computer Engineering at Georgia Institute of Technology; ELD is in the Department of Biomedical Engineering and affiliated with the School of Electrical & Computer Engineering at Georgia Institute of Technology

## Abstract

Models of neural architecture and organization are critical for the study of disease, aging, and development. Unfortunately, automating the process of building maps of microarchitectural differences both within and across brains still remains a challenge. In this paper, we present a way to build data-driven representations of brain structure using deep learning. With this model we can build meaningful representations of brain structure within an area, learn how different areas are related to one another anatomically, and use this model to discover new regions of interest within a sample that share similar characteristics in terms of their anatomical composition. We start by training a deep convolutional neural network to predict the brain area that it is in, using only small snapshots of its immediate surroundings. By requiring that the network learn to discriminate brain areas from these local views, it learns a rich representation of the underlying anatomical features that allow it to distinguish different brain areas. Once we have the trained network, we open up the black box, extract features from its last hidden layer, and then factorize them. After forming a low-dimensional factorization of the network’s representations, we find that the learned factors and their embeddings can be used to further resolve biologically meaningful subdivisions within brain regions (e.g., laminar divisions and barrels in somatosensory cortex). These findings speak to the potential use of neural networks to learn meaningful features for modeling neural architecture, and discovering new patterns in brain anatomy directly from images.

## 1 Introduction

Mapping out the underlying microstructure of the brain is essential for many tasks in neuroscience [1, 2, 3]. In studies of disease [4, 5], aging [6], or development [7], for instance, a rich description of the microstructure is first needed before being able to compare brains across different conditions. Detailed maps of brain structure have also led to important discoveries regarding relationships between structure and function [8, 9], and provide a necessary sign post when targeting specific brain regions for subsequent studies.

Much of our study of the brain and it’s organization, which dates back to Cajal’s beautiful renderings of neural architecture [10], still relies on human reasoning to define regions-of-interest (ROIs). This is done by examining different parts of an image, either in terms of their anatomical compositions or patterning, and then building an atlas or model of how the architecture changes across different brain regions. Moving forward however, given the ever-increasing sizes of new neuroimaging datasets, we need automated solutions that can find characteristics of brain structure, discover architectural patterns or primitives that are characteristic of a brain area, and also provide good ways to discover substructures (like cortical lamina) within a known brain area.

Unfortunately, when attempting to translate expert human knowledge into an automated method for region discovery, it is often unclear how to define the necessary image properties to extract. Moreover, extracting these features can be difficult to automate, or they may not be robust to minor changes in data (e.g., illumination conditions, background intensity, or noise) [11, 12]. Features therefore typically need to be hand-crafted for each individual dataset and application, and problems with generalization are only further exacerbated by biological variability, as well as process variations introduced during sample preparation, imaging, or post-processing [13]. Consequently, defining meaningful features in an image that can be used to define brain areas, is a very difficult problem.

Convolutional neural networks (CNNs) are particularly well suited for this task, given that they are designed to build and learn hierarchical [14, 15, 16], textural [17] representations directly from raw (image) data, and have been shown to conclusively outperform classifiers trained on hand-crafted features. These approaches are now being routinely applied in modeling brain structure in macroscale (MR) datasets [18, 19], and more recently for high resolution data [20, 21], to solve a variety of problems, ranging from tumor detection to pixel-level semantic segmentation in connectomics. Applications of deep learning to image understanding in microscopy, have so far focused on supervised approaches, where labeled data exists and the network is trained to segment data: either into brain regions [22, 23, 24], or into individual pixel-level components like neurites [25, 26]. While all of these approaches are capable of learning rich features from images to solve their respective tasks, they do not provide a tangible way to discover new areas or regions of interest, as they are designed to find specific components that they have been trained to segment.

In this paper, we introduce a deep learning-based approach for modeling microstructure in brain imagery (Figure 1, top). We envision that this approach can be used to automatically discover unique clusters or regions within a brain sample that share similar characteristics in their local morphology or cytoarchitecture. Our solution starts with the initial observation that if we train a network to do well at a brain area classification task, using only *local* views of the brain’s structure, then this forces the network to pay attention to specific anatomical features of the dataset and build rich feature sets (e.g., the patterning of axons, or the density and morphology of cells) along the way. After training a network to solve this task, we then open it up and peer into its hidden layers. We use the activations of units in the network’s layers as the “representations” of each input, thus treating the trained network as a feature extractor [27, 28, 29]. Because these features have been generated by a network trained to provide meaningful information about the diverse brain structures it has previously seen, they can potentially provide cues about changes or divisions within areas (e.g., layers and sub-divisions), and understand how different brain areas are related to one another.

**Figure 1:**
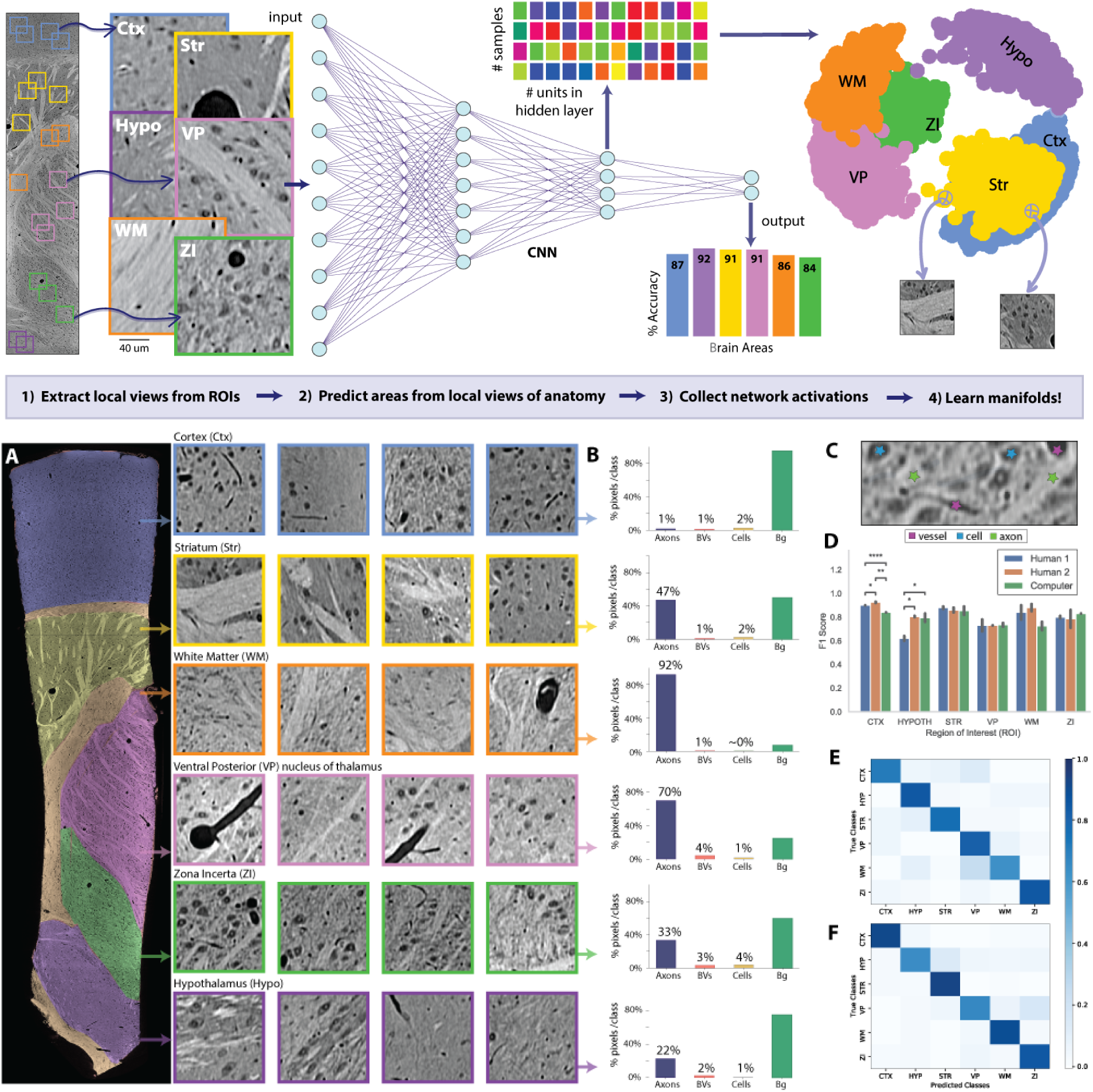
Deep feature learning approach for modeling brain microarchitecture. At the top, we provide an overview of our approach for neuroanatomical discovery. First, we select patches representative of different ROIs from a large brain sample, and use these examples to train a deep CNN that can classify local snapshots of the brain into different ROIs. We then extract activations from the trained network for many test samples and embed them into a low-dimensional space. Below, in (A), to the left we show an image slice from our dataset that spans six different brain areas and to the right we show exemplary image patches extracted from different areas that were used for the local classification task. Using the outputs of a network trained to perform semantic segmentation, in (B), we estimate the fraction of pixels in the image patches belonging to the different anatomical structures present, viz. axons, blood vessels, cells and tissue, visible in the X-ray image at this resolution. In (C), pixel-level components (vessels, cells, axons) are visualized and highlighted in an image from ZI. In (D), we quantify human annotator performance of inferring brain area from local snapshots like those shown in (A), by asking them to perform the local area identification task across a total of 360 images (60 per brain area). Their results were compared with those of a trained neural network (computer), and we also provide the statistical significance (independent t-tests) of the resulting analyses. Error bars used are standard deviations. (Note - * : 0.01 <= p < 0.05, ** : 0.005 <= p < 0.01, *** : 0.001 <= p < 0.005, **** : p < 0.001). In (E) and (F) we show the confusion matrices for the predictions of the trained neural network and humans respectively.

We applied this framework for microstructure discovery to a large-scale brain sample [30] imaged with synchrotron X-ray microtomography (micro-CT) [31]. This sample consists of six different brain areas that exhibit heterogeneous microarchitecture in terms of white matter as well as cell morphology and density. We trained a CNN to predict brain areas from snapshots of the local microarchitecture (150×150 micron image patches) within a larger image slice, sampled uniformly across different ROIs. Post training, we applied two dimensionality reduction techniques - principal component analysis (PCA) and non-negative matrix factorization (NMF) [32] - to the network activations generated from image patches drawn from an entire (test) brain slice (5.5M images). Both PCA and NMF provide factorizations of network activations that reveal biologically relevant information across the different brain areas and a lens into how the areas are organized within the network.

Further analysis of the network representations using NMF revealed that the non-negativity constraint is sufficient to generate *sparse* and *localized* factorizations. In this case, the embeddings of images onto these factors reveals different groupings or clusters within and across brain regions. A deeper investigation into these sub-regions reveals that different NMF components exhibit similarities in terms of their cell densities or local axonal projection patterns. Thus, by coupling this deep feature extraction method with a Gaussian mixture model (GMM) for clustering, both laminar differences and barrel fields [33] in cortex can be pulled out from the images without knowledge of these motifs. Thus, these findings point to the fact that deep learning-based representations can be used to find finer sub-divisions and biological features in the data, even when they’re not trained explicitly to do so. These advances can be translated into approaches for disease diagnosis, to model continuous variability in brain structure, and to discover micro-architectural motifs in new areas.

## 2 Results

In order to build an expressive model of local microarchitecture across heterogeneous brain areas, it was imperative that we first selected image data that has diverse microarchitecture, and was of sufficiently high resolution. To achieve this aim, we used a publicly available 3D X-ray microtomography dataset [30] (http://bossdb.org/project/prasad2020) that contains a diverse set of neuroanatomical structures, including myelinated axons, blood vessels, and cell bodies resolved at 1.17 micron isotropic (Fig. 1A). This sample, the Agmon-Connors slice [34], spans multiple diverse brain areas, including: somatosensory cortex (CTX) (barrel fields), striatum (STR), the ventral posterior (VP) region of thalamus, hypothalamus (HYP), and zona incerta (ZI). Each image in the dataset (1420×5805 pixels) contains examples from all brain areas - and in total across 700 images, spans a volume of 5 cubic mm. Thus this dataset provided us with sufficient heterogeneity, and high enough resolution, for our purposes.

From the raw data, we generated a collection of 150×150 micron images from the six manually annotated ROIs in the dataset. We used these data to train, validate and test a deep CNN on a six-way brain area classification task (see Methods, Figure S1 for details on the architecture. Our training set consisted of ∼2,000 images per class, while the validation and test sets contained 1,000 images per class, respectively. To ensure that our CNN was trained on data that were truly representative of the distributions of the different brain areas, when curating the training set, we sampled image patches such that they were contained within a single area, and also avoided patches that contained the background (outside of the brain sample). We validated and tested the network using images drawn from brain slices 50 and 100 microns away from the training image, respectively; however, when curating the validation and test subsets, we didn’t enforce the same restrictions on the sampled images as the training set. These data provided the necessary scale and resolution to capture anatomical differences across all the brain areas of interest.

The trained network performed well on both the validation and test datasets, thus providing evidence that the classifier could generalize to new images (Figure 2). The resulting classification performance (in terms of accuracy) was 100%, 89.77%, and 88.88% on the training, validation, and test datasets, respectively. Much of the degredation in performance can be attributed to the fact that the validation and test data were not as carefully curated as the training set, and many images in these sets also spanned boundaries between areas or lay close to the edge of the sample. When viewing images at the same scale, human experts classified a subset of the image patches (60 per class) drawn from the test slice (z = 259) into areas with ∼ 81% accuracy, and with similar types of errors overall (Figure 1D-E) [30, 35]. In general, cortex and striatum appear easiest to classify, and hypothalamus, due to its diversity and similarity to other areas, is hardest to classify correctly. Due to similarity in cell types and morphology between VP, hypothalamus, and ZI, these areas can also be easily distinguished from cells in cortex and striatum. Critically, we find that the visual cues that one needs to use to determine which area you are in at this scale requires focusing on specific anatomical structures like axons, as well as the cell morphology and density in the image.

**Figure 2:**
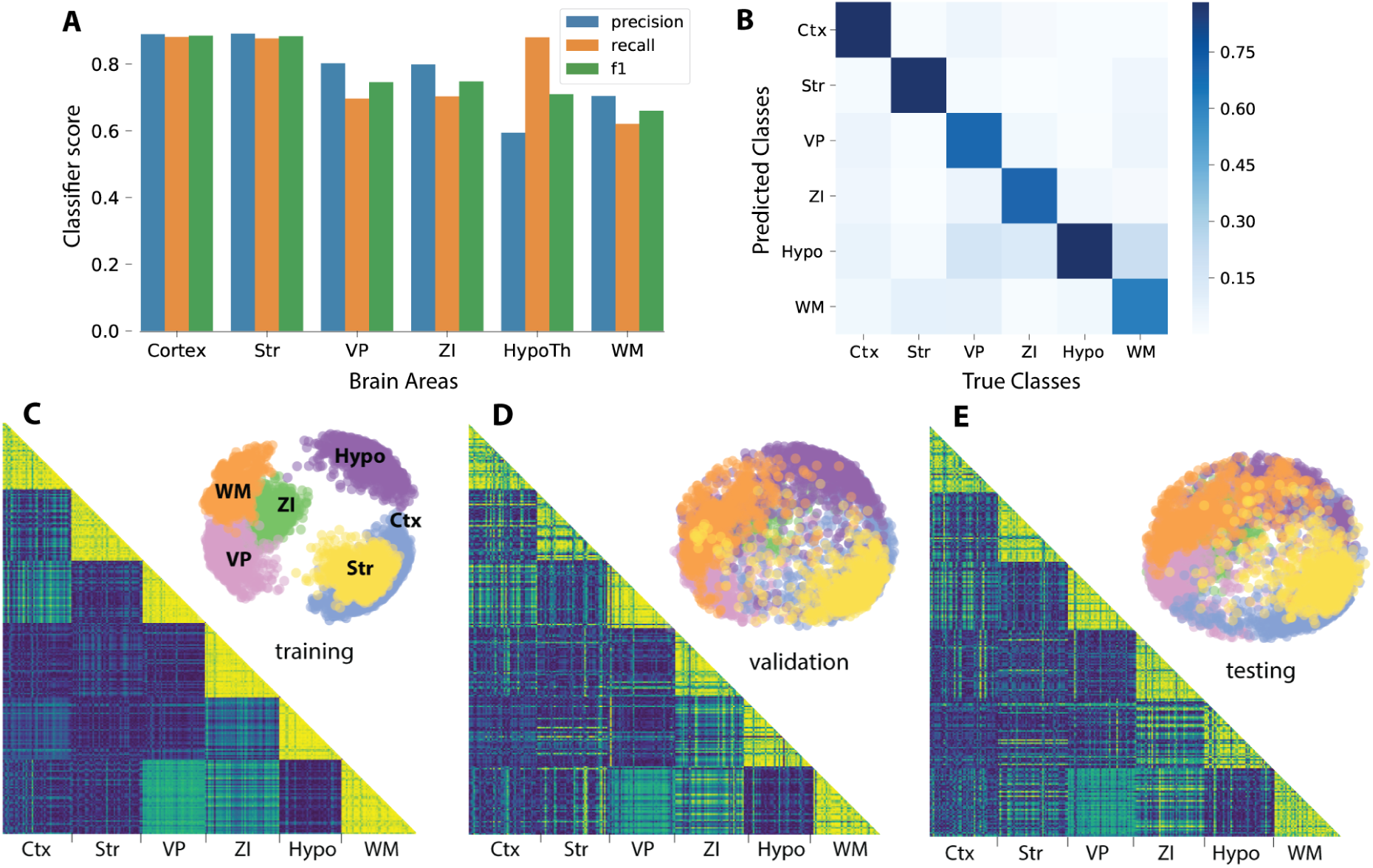
Evaluating network performance across training, validation and test data. In (A) we show the precision, recall, and f1 scores for the trained network across areas and in (B) we show the confusion matrix formed using the network’s predictions on the test data. Both of these figures were generated by running the network at scale on the entire test slice (z=259). In (C), (D) and (E) we show the covariance matrices computed from the network representations collected for the train, validation, and test sets, respectively. These matrices are symmetric and thus only the bottom half of the matrix is shown. In their place, we show the 3D embeddings of the activations obtained via PCA.

To further examine the generalization performance of the network, we densely sampled points in the test thalamocortical slice (z=259) and predicted the class of the image immediately surrounding each pixel, resulting in ∼ 5.5M images for quantification. Over this set, we obtained performance between 85% (cortex, striatum) to 55% in hypothalamus (Figure 2A-B). These results appear to align with our previous smaller scale estimates in terms of which classes are easiest or hardest to classify. By further enforcing local consensus amongst the class estimates, the pixel-level outputs of the CNN can be denoised through a simple nearest neighbor post-processing step (see Algorithm 1 in Methods; Supp. Materials S2). After this post-processing, the performance on a full thalamocortical section (from cortex to hypothalamus) increases to over 90%. Thus we show that it is indeed possible to train a deep learning model to discriminate brain areas from local snapshots of anatomy at scale.

After training the network to discriminate brain areas, we then asked whether it was possible to use the same network to explore the finer-scale microarchitectural characteristics within the input images. To do so, we used the network in a fashion similar to how pre-trained networks are used in the machine learning community [27, 28, 36, 29]. In these settings, an input is passed through a trained network (with its weights fixed) and then the activations of units at specific points in the network can be read out and interpreted as a feature (Figure 1). Using this approach, we froze the weights and passed samples through the network to generate a set of multi-dimensional (64D) network activations by pulling from the last hidden layer, over image patches across all six brain areas. We computed and visualized the covariance of these network activations across the training, validation, and test sets (Figure 2 C,D,E), and found that in all cases, the trained CNN network did a good job of pulling apart the classes that it had been trained to distinguish, as evidenced by the class-wise block structure in the covariance matrices. When analyzing the representations, we found that areas that are far from each other in terms of their anatomy (VP vs. CTX) remained well separated even in the representation space of the network. We found that even though many of the images collected from striatum and cortex were very different in terms of their pixel-level content (i.e., striatum has an abundance of axon bundles, cortex has very few myelinated axons that are visible), the network learned features that kept the two classes close to one another in the latent space, which is also a strategy used in human experts performing the same task. The images from white matter tracts and the VP region of thalamus are also organized close to one another in the latent space, again aligning with human intuition, given the presence of heavy amounts of patterned WM in thalamus in this section. Thus, features extracted from the trained CNN provide meaningful information and preserves relationships amongst classes.

We next conducted a large scale experiment and extracted the network activations for approximately 5.5 million image patches that densely tiled an entire virtual section, (test image slice, z=259) spanning cortex all the way down to hypothalamus (Figure 3). We then computed the principal components (PCs) of the network activations for these images, and projected all of them into this learned low-dimensional space (Figure 3B). By stacking the first three PCs into the different color channels, we could visualize the micro-architectural variability across all three PCs simultaneously. At scale, the first three principal components together are almost sufficient to distinguish the different brain areas as the PCs map to unique colors. This is further confirmed by clustering the image patches according to their embedding along the first three PCs (Figure 3C, ARI = 0.703). When we examined the representations across each PC, we found that multiple brain regions are highlighted within the same component, and found similar overall structural findings as before in the small-scale datasets (e.g., cortex and striatum are encoded in the same PC). We also visualized a few more components and found results along similar lines for all the visualized PC embeddings (Supp. FIgure S4). These results point to the fact that the features formed within the network provide a compact low-dimensional embedding of the data when viewed at this macroscale.

**Figure 3:**
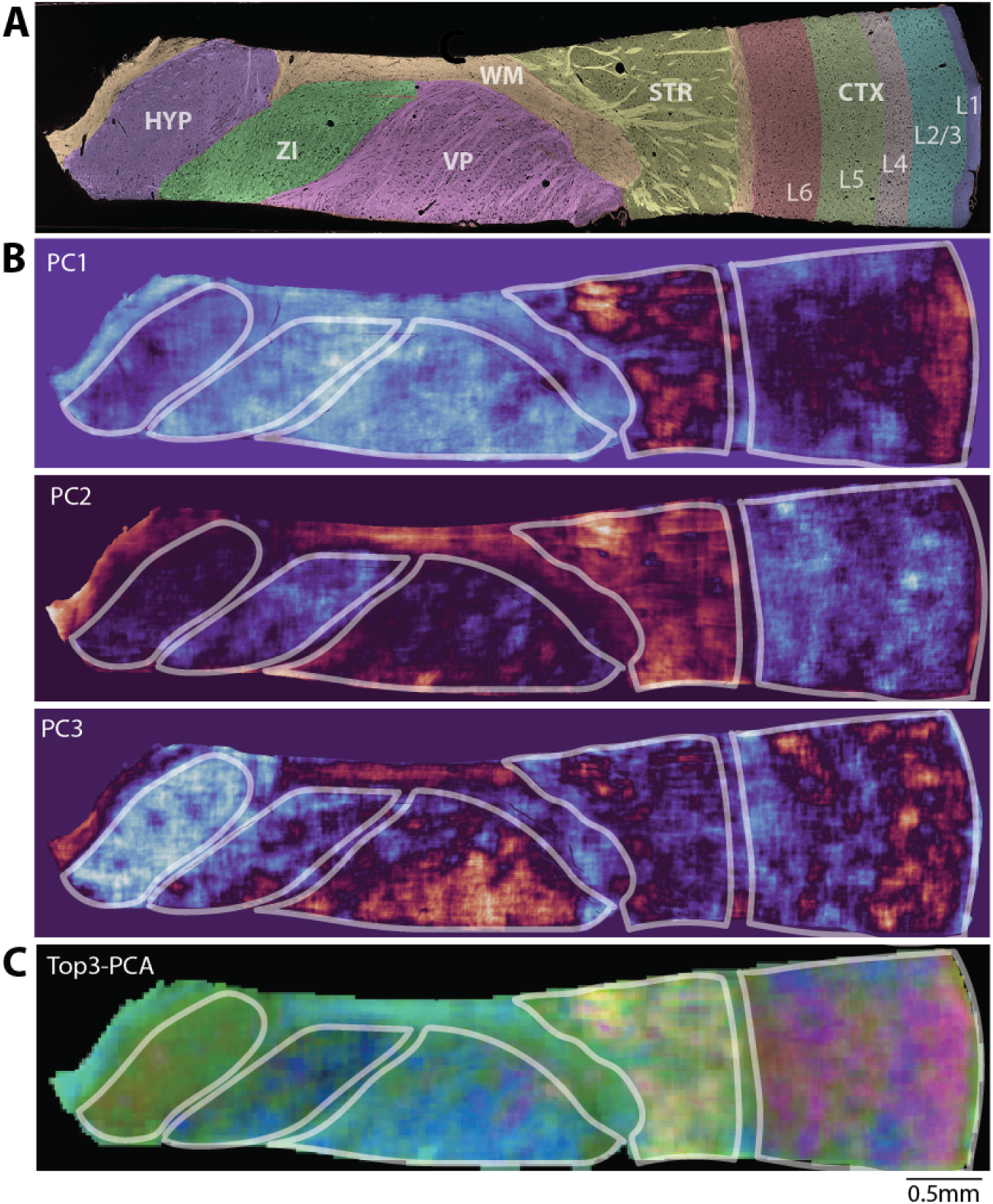
Visualization of low-dimensional PC embeddings of network activations. In (A) we show the manual annotations for all brain areas in the sample (z=259). The cortex is further divided into its different layers. In (B), we visualize the low dimensional embeddings along the first three PCs. (B-PC1) shows particularly pronounced positive expression in axon-rich parts of the striatum, along with layer 1 of the cortex (regions that are more orange/red) and negative expression in thalamic regions (white/blue regions). High positive expression can be interpreted as a strong positive correlation of the embeddings with the PC while high negative expression reflects strong anti-correlation. (B-PC2) reveals axon-rich areas in general while (B-PC3) shows high negative expression in the hypothalamus and positive expression in some axon rich regions, particularly the VP. We visualize the PCs together in a multi-channel image in (C), where the red, green and blue channels show the correlation of the embeddings with the first, second and third PCs respectively.

Although PCA enabled an initial visualization of how brain areas are related to one another, we asked whether imposing additional structure in the factorization could provide further improvements. Thus, we applied an alternative technique for dimensionality reduction, called non-negative matrix factorization (NMF). NMF constrains all components of the factorization to be non-negative, and critically, it also relaxes the assumption that they need to be orthogonal. These properties of NMF allow it to pull out factorizations that have some overlap between each other in the low-dimensional space (unlike PCA) and also endow it with a certain ability to inherently cluster samples. Consequently, NMF revealed sparse, localized expression within a number of its factors (Figure 4, Supp. Materials Figures S5-S10), and interpretable co-expression across multiple ROIs, as opposed to the relatively more dense expression that we see in the PCs.

**Figure 4:**
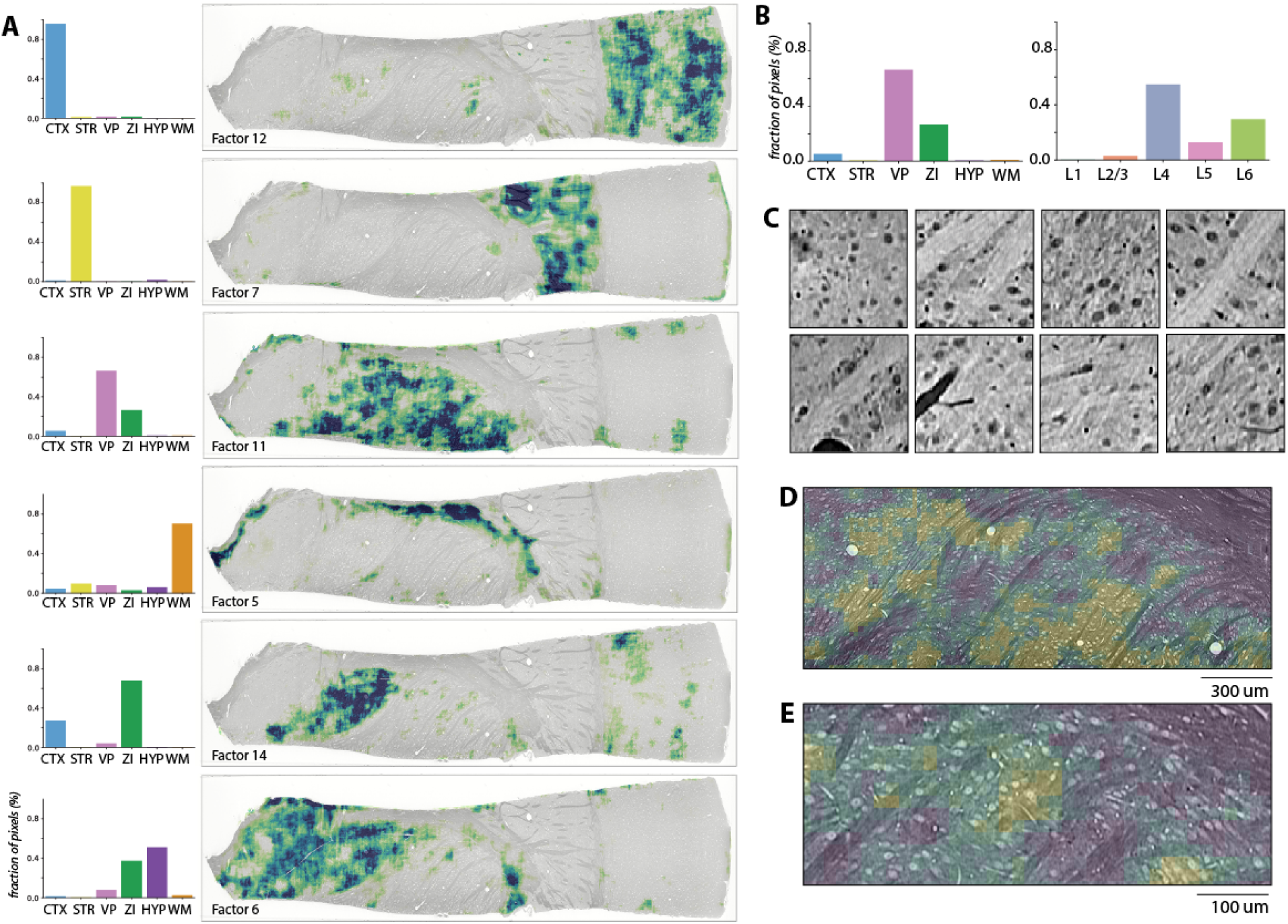
Non-negative matrix factorization applied to an entire image slice from a thalamocortical section spanning six brain areas. We show embeddings of the extracted CNN feature representations along a subset of the top fifteen factors that were obtained using non-negative matrix factorization (NMF) on the entire image slice. In (A), we analyze the coefficients of these factors and draw the following conclusions: Factor 12 (F12) highlights most of cortex (likely areas not heavily populated with axons). In F11, the regions of VP that are densely populated with axons and parts of white matter are highlighted. Factor 7 (F7) appears to be heavily aligned with Striatum while F6 aligns itself with most of the hypothalamus and axons of the striatum, alongwith white matter tracts. F5 almost exclusively highlights white matter. The barplots associated with each of the factors show the distribution of signal in each of the different brain areas, for the respective factors. To the left of the non-negative embeddings, we also show the distribution of the signal for the chosen factors across the different brain areas. The distribution of expression in F11 across the different brain areas is seen in (B-left) while the distribution of expression in different layers of the cortex is shown in (B-right). In (C) we show the 8 images that most strongly align with F11. It is easy to see that the factors localize particular brain areas, with F12, F7 highlighting the cortex and striatum respectively, F11 highlighting thalamic regions (VP and ZI) and F14 resolving only the ZI. With (D-E) we zoom into the expression of F11 in parts of the thalamus that reveal that this factor aligns with patches which simultaneously contain both axonal matter and cells.

Since the factors provided by NMF are unordered in terms of how much of the variance they explain in a dataset, we needed a way to visualize factors that may highlight biologically meaningful structures in data. Thus, we selected six factors that are aligned with the six labeled brain areas in the dataset and yet still uncorrelated with one another (see *Methods - Selecting a subset of predictive factors*). Factors F12, F7 and F14 are all selected as factors that are most aligned with the cortex, striatum and ZI, respectively, while F11 and F5, F6 reveal co-expression patterns that align with specific neuroanatomical features (e.g. regions with myelinated axons with little to no cells for F5 and a general thalamic distribution in F11). Further examination of Factor 11’s embedding highlights parts of the image (Figure 4C) that have patchy and diffuse axonal expression, which mainly highlights VP and boundaries of the WM tracts, along with some parts of ZI innervated with axons. However, unlike other components that highlight WM specifically (F5), this factor also seems to require some distribution of cells as well (Figure 4D, E). Studying the signal distribution of F11 (Figure 4B) reveals that the factor shows activity in the VP, ZI and to some degree, the cortex as well. While it isn’t startling to see co-expression of regions in the thalamus, to see the same factor light up parts of cortex was surprising. When we further zoomed into F11’s expression in the cortex, and examined the component correlation across manually annotated layers, we observed that this same factor also highlights two distinct parts of both Layer 4 (barrel fields, somatosensory cortex) and Layer 6, both of which do indeed exhibit diffuse expression of axons (Figure 4B-right). Picking out these regions of sparse axonal innervation by eye is not easy and the network appears to have identified a good solution to this problem. This analysis therefore provided us with insights into how different brain areas are micro-architecturally related, and how patterning of axons and cells appear to be aligned with specific factors.

Our analysis also revealed interesting substructures in cortex that appeared to map roughly onto different cortical layers (see Supp. Materials Figure S12,S13). Further subdivisions in Layer 4 were noticeable, which appeared to alternate in higher and lower density regions; these patterns of architecture align with descriptions of the patterning of barrels and barreloids in somatosensory cortex (Figure 4A) [33]. This patterning was further evident when zooming into the top three NMF components selected to separate the underlying cortical layers (see Methods). We then used the top 15 factors obtained with NMF to cluster data by fitting a Gaussian Mixture Model (GMM) (see *Methods - Gaussian mixture model fitting and clustering*), for both the test slice (z = 259, Figure 5C) and another slice 100 slices away (z = 359, Figure 5D). In both cases, we find similar distributions of coefficients (embeddings) across the four clusters (mixture components) that also align nicely with laminar boundaries (Figure 5E). We observe that two components represent images that have a reasonable cell density and have no axons, with Layers 2/3 and 5/6 being split roughly across the two. Examining these components further revealed that they appear to be aligned with slight differences in cell density. Consistently, parts of the image that have high axon count and also high cell density get labeled and grouped into one component, both in Layer 4 and 6. Inspection of the raw image data at higher resolution (Figure 5F-G) confirmed that the patterns observed in NMF embeddings, align with laminar boundaries, clusters of neurons that align with the barrel fields in Layer 4 and barreloids (regions of low density between barrels). These results provide exciting new directions for using dimensionality reduction to get a view into neural network representations, and learn how different parts of the sample relate to one another.

**Figure 5:**
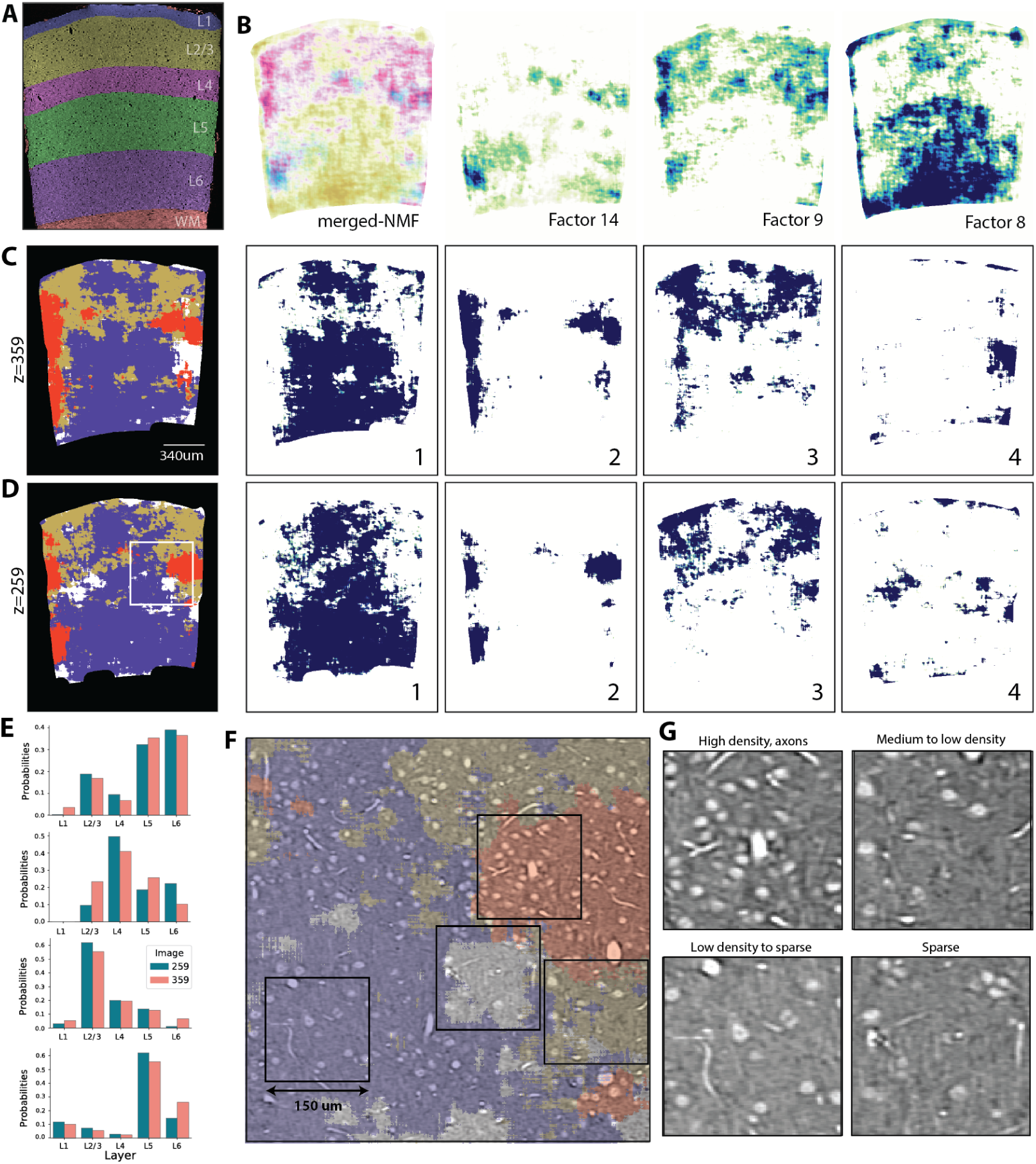
Clustering representations in cortex to reveal laminar divisions and regions of varying cell density. In (A), we show the annotations of cortical layers in the barrel field region of somatosensory cortex. Five distinct layers can be resolved in this region on the basis of cell size and density, with layers 2 and 3 being grouped into one in our dataset. In (B), we show the cortical embeddings for three factors that are selected to be uncorrelated across the layers while still spanning most of the cortex. To the left of the three factor embeddings, we show a merged color representation, with each of the three factors mapped to a unique color and overlaid (F14 is cyan, F9 is magenta, and F8 is yellow). In (C-D), we show the GMM clustering results for two different test slices (z=259, z=359). For both, we have the clustering results using a GMM (*n*_*g*_ = 4) on the very left. For both slices, we used the distribution of signal across the layers to match the two factors to one another. We also observe that all three of the chosen factors we see in (B) are revealed in the clustering results when applied to the 15D NMF representations. In (E), we show the layer-specific expression for each factor after normalizing it into a distribution, for both slices. These results show strong similarity between the factorizations of the two slices, and localization across layers in many cases. We zoom into a region of cortex in (F-G), and examine different high and low density regions detected after clustering.

## 3 Discussion

In this work, we presented a deep learning strategy for modeling brain structure and organization. We showed that by training a neural network to classify brain areas from only local snapshots of brain structure, and then decomposing the representations formed by this network, it is possible to reveal interpretable differences between brain areas and find finer-scale substructures within the sample (e.g., low or high cell or axon density regions/clusters in cortex). We showed that this general framework, where dimensionality reduction are deep feature extraction are married, can be used to discover biologically relevant sub-divisions and variations within and across brain areas with diverse cytoarchitecture.

When designing the network, we selected patches that are 150×150 microns in size; with this field-of-view, the resulting images were small enough to limit global context (e.g., the relative positioning of different brain areas) but also large enough to span multiple cells and thus give an estimate of density, and a view into the local patterning of axons. Since we do not operate on the entire image at a time, but on small, local patches extracted from the image slice, the network cannot use the global structure of the areas in the entire slice (including background information), but must instead truly learn the local distributions of different brain areas. As the scale will alter the types of patterns that the network pulls out, it would be interesting to examine how different scales of images impacts the performance and what the network learns to group, and how this approach may even be extended to incorporate information across multiple scales.

Another important design decision that was necessary to consider was *where* in a network we should target for our analysis. An intuitive reason why we pull representations from the last hidden layer, rather than those from previous layers, is that as we go deeper into the network, the features in a layer grow increasingly task-specific [37]. While representations from the first few layers contain the most information about the raw input image itself [38], representations at later layers form efficient codes [39] that encode information not only about the image, but also about the class and the features in the image that distinguish it from those belonging to other classes. In situations where one would want more “general” features that aren’t task specific (e.g., transfer learning between a source and target domain that aren’t similar) - extracting features from the lower layers of the networks and appending a few layers on top of the extracted features and training them, might provide a more meaningful result. Exploring the representations formed across different layers in the network, and examining how they can be used to build models of brain structure and organization is an interesting direction for future research.

In the past few years, the use of ImageNet pre-trained networks for feature extraction and transfer learning in biological data has grown immensely [22, 40, 41]. However, the effects of pre-training on *natural images* for transfer learning with medical data are, at best, not completely understood or at worst, harmful, since the underlying distributions of the two types of data are significantly different [42]. While pre-trained networks provide a way to quickly transfer previous training data to a new task and allow for the construction of good one-vs-rest classifiers to separate data, our previous work [43] shows that these representations may not be stable. Because these representations lack separability in terms of the desired class structure, while a classifier can be trained to map inputs to the desired class, the introduction of even small amounts of noise can move the representation far from the true class. In comparison, when a network is trained to solve a brain-area discrimination task, the resulting representations exhibit high degrees of separation. These findings, both in terms of the lack of separability and robustness of representations generated by pre-trained networks, suggest that classification performance is not sufficient to gauge whether a representation is good for a downstream task. When going beyond classification, and aiming to perform the sorts of unsupervised analyses that we introduced here, well organized and structured representations to compare and contrast images, are of paramount importance.

This work shows how knowledge of the latent space of network activations can provide a way to map further sub-divisions in tissue that the network hasn’t necessarily been trained to detect. Here, we applied a mixture model to identify these divisions or clusters within an area; one could also apply hierarchical clustering methods to build a flexible decomposition of a sample into grosser areas first and then subdividing within at a finer spatial scale. Additionally, the learned features could also be used to find anomalies or deviations in the representations formed over standard and healthy brains to determine when an input image is an outlier or might be pathological. The representational learning strategies presented here therefore provide a path forward in discovering patterns in brain architecture without the need for labels, and provides a principled way to leverage deep learning in comparative neuroanatomy and digital pathology.

Traditional methods for brain mapping assume that there is a common atlas and thus there are common boundaries that separate distinct areas [44, 45]. However, we know that borders are not always sharp and can instead be stochastic and vary across individuals [22, 46], and hence on average can be best modeled with probabilistic transitions. With the methods that we’ve introduced, we can compare and contrast the cytoarchitecture of brains in exquisite detail, thus revealing maps that can be compared and contrasted across individuals to build not only probabilistic models for boundaries, but also a multi-dimensional descriptions of how the brain’s structure shifts as we move from one area to the next. By viewing brain structure and architecture in terms of the many dimensions over which it can vary, we will be able to build improved models to understand how individuals differ, and how the brain is impacted by disease, learning, and development.

## Methods

### Convolutional neural network architecture and training

We trained a deep convolutional neural network on a brain area classification task, where the goal was to determine whether a 128×128 image in the dataset was drawn from: Cortex, Hypothalamus, Striatum, the Ventral Posterior nucleus of thalamus, White Matter (corpus callosum, internal capsule), or Zona Incerta. Our CNN had four convolutional layers with kernel sizes 7×7, 5×5, 3×3 and 3×3 respectively, followed by three fully connected layers with 1024, 64, and 6 nodes at the output. We also used dropout (*p* = 0.3) between the fully connected layers of the network (see Supp. Materials S1 for a visualization of the architecture and further details). The activation function used throughout was the rectified linear unit (ReLU) and the loss function was regular crossentropy. The feed-forward network was trained over 100 epochs using the Adam optimizer with a learning rate of 1e-4. The best model was chosen on the basis of accuracy on the validation set. All aspects of the CNN modeling and training were carried out in PyTorch [47] and Torchvision. The model and weights are available for download [48].

### Nearest-neighbors cleanup for large-scale brain area segmentation

Once we had the trained model, we deployed it on a large image spanning all six brain areas to test the network’s ability to reliably identify different brain areas at scale. Even when the CNN gets most of its predictions right, there were many small clusters throughout the image that had been misclassified. We therefore developed a K-Nearest Neighbours (kNN) based cleanup procedure (Algorithm 1) that enforced local spatial consensus amongst patch predictions. The algorithm correctly re-mapped many small clusters of misclassified patches to the correct area, thus significantly boosting the overall accuracy of the resulting area-level segmentation (Supp. Figure S2). The main idea behind the algorithm is simple: if the *i*^th^ ROI is expected to have *b*_*i*_ spatially contiguous parts (connected components), then we can keep the labels of the points in the *b*_*i*_ largest components that have been predicted as belonging to the *i*^th^ class, and “remove” the labels of the points that didn’t belong to the *b*_*i*_ largest components. After repeating this step for all *C* classes, all samples without labels are then re-labelled with the label of the majority of their *k* spatially nearest neighbours that have labels.

In our implementation, we set *k* = 3. The features that we used when performing kNN and predicting the classes were the coordinates of the points in the full image, and the distance metric used was the Euclidean norm.

#### Algorithm 1 Nearest Neighbor Cleanup Algorithm

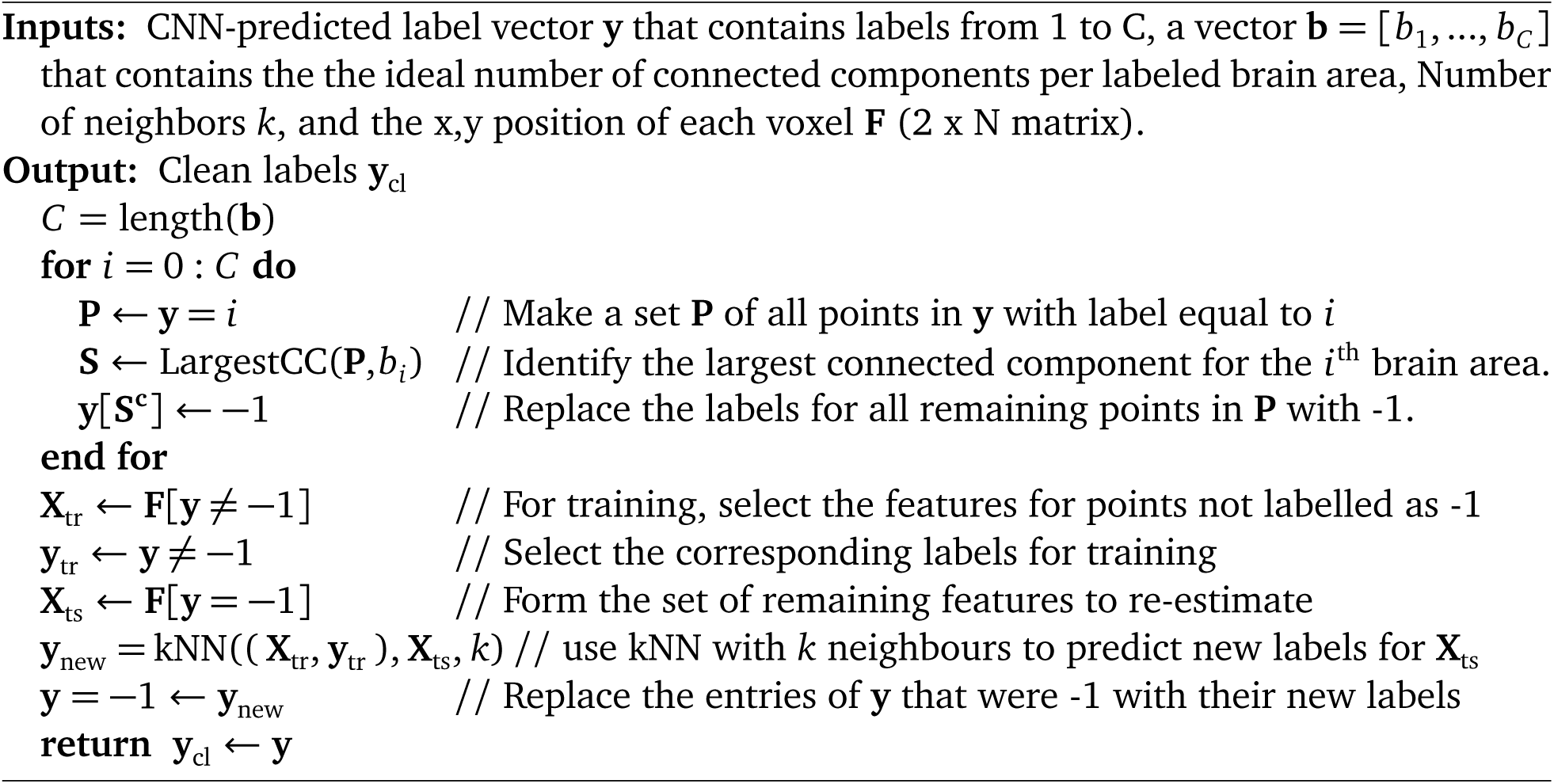

### Extracting network activations after training

After training the model, we passed a set of images **D** through the network and collected the activations from the last hidden layer. We then arranged these representations into a (*d, n*) matrix **X**, where **X** = *f*_*θ*_ (**D**), and *f*_*θ*_ (·) denotes the transformation from the original images to the last hidden layer computed by the network, and is parameterized by *θ*. In our experiments, *d* = 64 and *n* = 11080*/*6000*/*6000 for our training, validation and test datasets respectively. When computing the class-wise covariance (Figure 2), we first normalize the representations to each have unit Euclidean norm, and use the un-normalized representations in the rest of our analyses. We provide code to extract representations from our model using PyTorch at [48].

### Decomposing network representations using matrix factorization

The general matrix factorization framework is simply to approximate a matrix **X** ∈ ℝ ^*d*×*n*^ using two (or three) factor matrices **U** and **V** such that the residual ‖**X−UV**‖ is minimized with respect to a chosen metric, subject to a set of user specified constraints. Here, the matrix **U**∈ ℝ ^*d*×*k*^ is a basis set that projects the *d*-dimensional data onto a lower *k*-dimensional space and **V**∈ ℝ ^*k*×*n*^ is the matrix of coefficients obtained for all *n* examples. Each of the columns of **V** explain how aligned each data sample in **X** is to the different basis vectors that are given by the columns of **U**. What different matrix factorization methods differ in are the constraints that they impose on their factors, thereby resulting in different decompositions of the same data matrix **X**. The two factorizations that we used were the PCA and NMF, which differ from each other in the following key ways: i) PCA requires the columns in **U** to be orthonormal, while NMF does not require orthogonality in its basis elements. ii) NMF requires that **U, V** ≥ 0 but PCA does not. Consequently, their resulting factorizations and embeddings have different properties. PCA captures the most variance in the data and has the least reconstruction error when restricted to a rank-*k* approximation, but its requirement of orthogonality amongst its basis elements can result in embeddings that aren’t very interpretable and counter-intuitive, as there often isn’t a reason to impose such a constraint on the factors when working with real data. NMF however replaces orthogonality on the basis set with non-negativity in its factors, which many times is a reasonable assumption to make (i.e. none of the basis elements can make negative contributions). Furthermore, it results in an inherent clustering on the columns (i.e. samples) of the data matrix **X**, thus forming sparse, localized embeddings [49] that are easy to interpret.

We formed our feature matrix **X** by collecting activations from the last hidden layer of our trained network. For our experiments with both PCA and NMF, we set *k* = 15. To visualize the alignment of image structures in **X** with the different learned low-dimensional factors, we create a “heatmap” which colors each pixel with its corresponding coefficient in the matrix **V**. We found that PCA did not reveal any interesting patterns in the embedded space beyond the sixth factor and therefore show PC-embeddings only up to *k* = 6 (Supp. Figure S4). When visualizing the PC-embeddings as a multi-channel image (Figure 3), we used the first three PCs, and stacked each of the PC coefficients into the R, G, and B channels. Experiments were carried out in Python using the scikit-learn library [50]. Associated code and data are also provided [48].

### Computing regional and layer-specific distributions

Once we computed the factors and their associated embeddings, we needed a way to quantify how aligned the different factors were with the different ROIs in the sample. We did so by measuring the strength of the signal across different ROIs for the different factors, i.e., we summed across the columns in **V** that corresponded with the specific ROIs. We defined our ROIs as different brain regions when working with the entire image slice, and different cortical layers when analyzing the cortex. We then computed ROI-specific distributions for each of the factors and scaled them such that the distribution across the sample for every factor summed to 1. In the case of the entrire brain sample, the distributions sum to 1 across brain areas, whereas when analyzing just the cortex, the normalized density is conditioned on being in the cortex, and the distributions across the different cortical layers sum to 1 for each of the different factors.

The annotations for the different brain areas and cortical layers were provided by a trained neuroanatomist for one in every ∼50 slices across the entire three-dimensional dataset. Additional details about the data and how the ROI annotations were curated can be found at [30]. The ROI annotations are available openly at [51].

### Selecting a subset of predictive factors

PCA has a deterministic way of finding factors that is agnostic to the initial condition or the rank of the required approximation i.e., the first, second, third, etc. principal components of a matrix **X** remain the same irrespective of whether you solve for a rank-3, 6, or 15 approximation. This follows directly from the fact that the first PC is the direction which captures maximum variance in the data, the second the next most in a direction orthogonal to the first, and so on. This property of the PCs also gives them a natural sense of ordering, in that PC-1 contributes most to the reconstruction of a rank-*k* approximation, followed by PC-2, 3 and so on. Unfortunately, NMF does not result in factors that are any more or less important in explaining or reconstructing the data by any inherent property of the data matrix **X**, thus making it difficult to select a subset of predictive factors.

We therefore developed a greedy algorithm to choose the top-*k* predictive factors from a pre-computed set of non-negative factors for the purposes of visualization and post-hoc analyses. Our goal was to select the subset of *k* factors that shared as little information with each other in the different ROIs while still being able to explain as much of the entire sample as possible. For example, when working with the cortex as the target area, one would want to choose *k* factors that cover as much of the cortex as possible, while sharing minimal information across the different cortical layers. To do so, the algorithm takes as inputs the entire set of non-negative factors, the number of factors that one wants to sub-select, annotations that demarcate the different regions of interest in the sample and a tunable hyperparameter *λ*. It then computes two sets of scores called the “coverage scores” and “leakage scores” for all combinations of the different non-negative factors and ROIs in the sample. For a factor *i* and an ROI *a* whose set of points are given by the set *A*, the coverage score

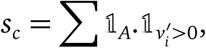

i.e., it is the number of positive coefficients for the ROI associated with factor *i*. Note that 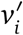 is *i*^th^ the column of **V**^**T**^. The leakage score for the same factor *i* and ROI *a* is

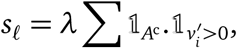

i.e., it is the number of positive coefficients outside the ROI and associated with factor *i*, scaled by *λ*. In case there are *f* factors and *r* ROIs, the coverage scores **S**_**c**_ and leakage scores **S**_**l**_ across all factors and ROIs would each be a matrix of size(*f, r*). The total scores across the different factors and ROIs is calculated as

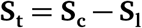

which again, results in a matrix of size (*f, r*). Once the total scores are calculated, the algorithm selects the factor that contains the cell with the single highest score and then the next one, given that it neither selects the same factor, nor another factor whose highest score is in the same ROI as the one selected previously. The procedure continues until the *k*-top scoring factors have been chosen.

In our experiments, we set *λ* = 0.5. Intuitively, larger values of *λ* would encourage sparser and more localized factors to be selected while lower values would allow for denser factors to be selected as signal outside of an ROI would not be penalized as heavily. We provide a Python notebook with the code for the method described above at [48].

### Gaussian mixture model fitting and clustering

We performed a clustering analysis on the 15-dimensional non-negative embeddings of representations from samples in the cortex from two different slices that were ∼100*µ*m apart (z = 259, 359), using a Gaussian mixture model (GMM). Prior to fitting GMMs on the cortical representations however, we got rid of the representations belonging to samples along the boundaries of the cortex so as to avoid outliers interfering with the analyses. The GMMs were fit to the representations from the two different slices independently using the expectation-maximization (EM) algorithm. However, even though the same number of Gaussians were fit to the representations across the two slices, clustering algorithms do not enable a direct correspondence between clusters predicted with the same label across different instances, i.e., the cluster that is labelled “1” in slice 259 would not necessarily correspond to the cluster labelled “1” in slice 359. To match the different clusters across the two slices, we therefore carried out the following procedure: i) We first computed the layer-wise cortical distributions (in the same way as described previously) for the different clusters. ii) We chose the order of components in one set of distributions as the “correct” order (In our case we took the order of components obtained for slice 259 as the correct ordering). iii) We then computed the Jensen-Shannon divergence between all the distributions for slice 259 with all the distributions in slice 359, thus giving us our distance matrix. Assuming that we fit a GMM with *n*_*g*_ components on the representations for each of the slices, this step would result in a matrix of dimensions (*n*_*g*_, *n*_*g*_). For each of the *n*_*g*_ clusters for slice 359, we match it with, i.e., assign the same label as the cluster in slice 259 that it has the least distributional distance with.

The number of Gaussians that we used in our GMMs was set to 4. All experimentation was conducted in Python with the Numpy and Scikit-learn libraries. Code for the conducted analyses is provided [48].

## Code and Data Availability

We provide the: (i) training, test, and validation datasets, (ii) pre-trained network architecture and weights (PyTorch), (iii) network activations for all three datasets, and (iv) Jupyter notebooks for computing and running our distributional and clustering analyses on the precomputed representations at https://nerdslab.github.io/deepbraindisco/.

## Author contributions

ELD and AHB conceived of and designed the research methods, and wrote the manuscript.

## Acknowledgements

We would like to acknowledge Judy Prasad for her insights and help in interpretation of the X-ray dataset, regional and cortical layer annotations, as well as helpful discussions and feedback throughout the project. We would also like to thank Julie Harris and Jennifer Whitesell for helpful conversations and feedback at different stages of this project. Thanks to Keren Zhang, Nischita Kaza, Namrata Nadagouda and Konrad Kording for helpful feedback on the manuscript. This work was supported by NSF award number IIS-1755871 (ELD, AHB); ELD was also supported by an Alfred P. Sloan Fellowship and NIH Award No. 1R24MH114799-01.

## Supplemental Materials

**Figure S1:**
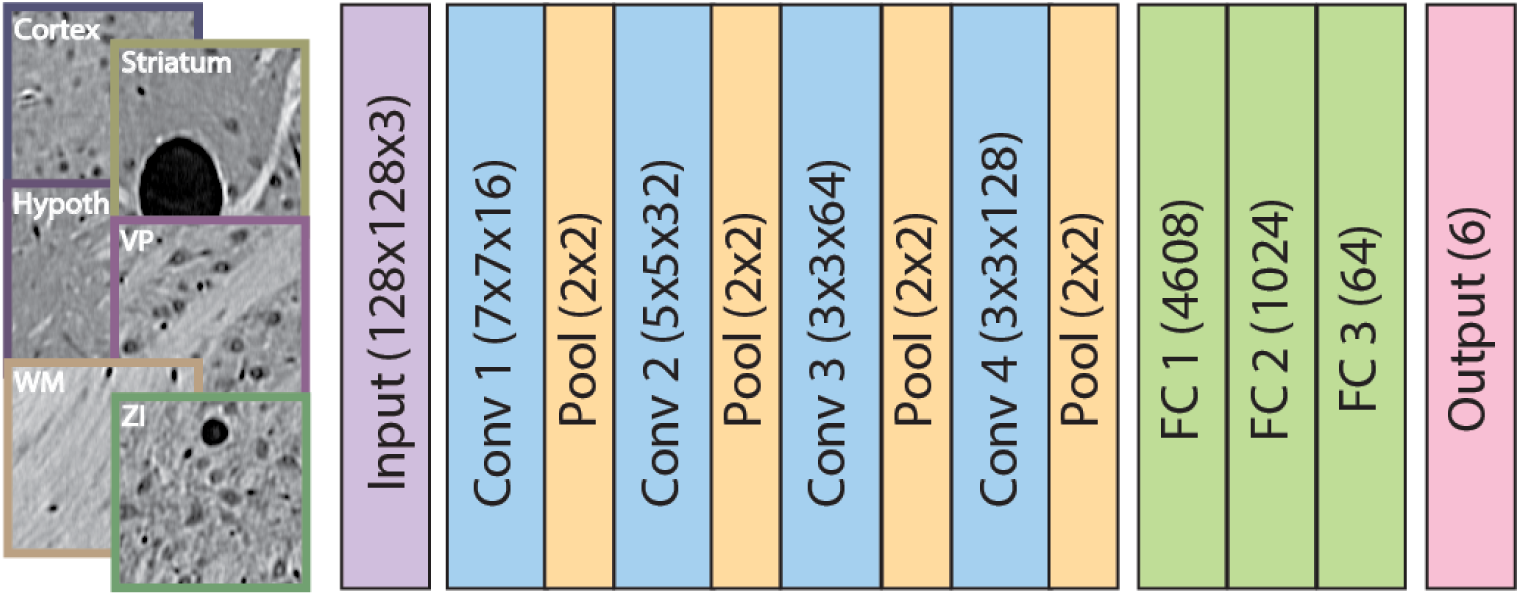
CNN architecture. The network used in our experiments is a 7-layer feed-forward deep convolutional neural network. The (*k, k, c*) convoltuional layers (shown in blue) have *c* filters in the layer, each with a receptive field of size (*k, k*). All the convolutional layers have stride = 1. The first two layers have padding = 1 while the last two convolutional layers don’t use any padding. The pooling layers (shown in yellow) used throughout perform max pooling and have a stride of 2. The fully connected layers (shown in green) and each of them has *d* nodes in the layer.

**Figure S2:**
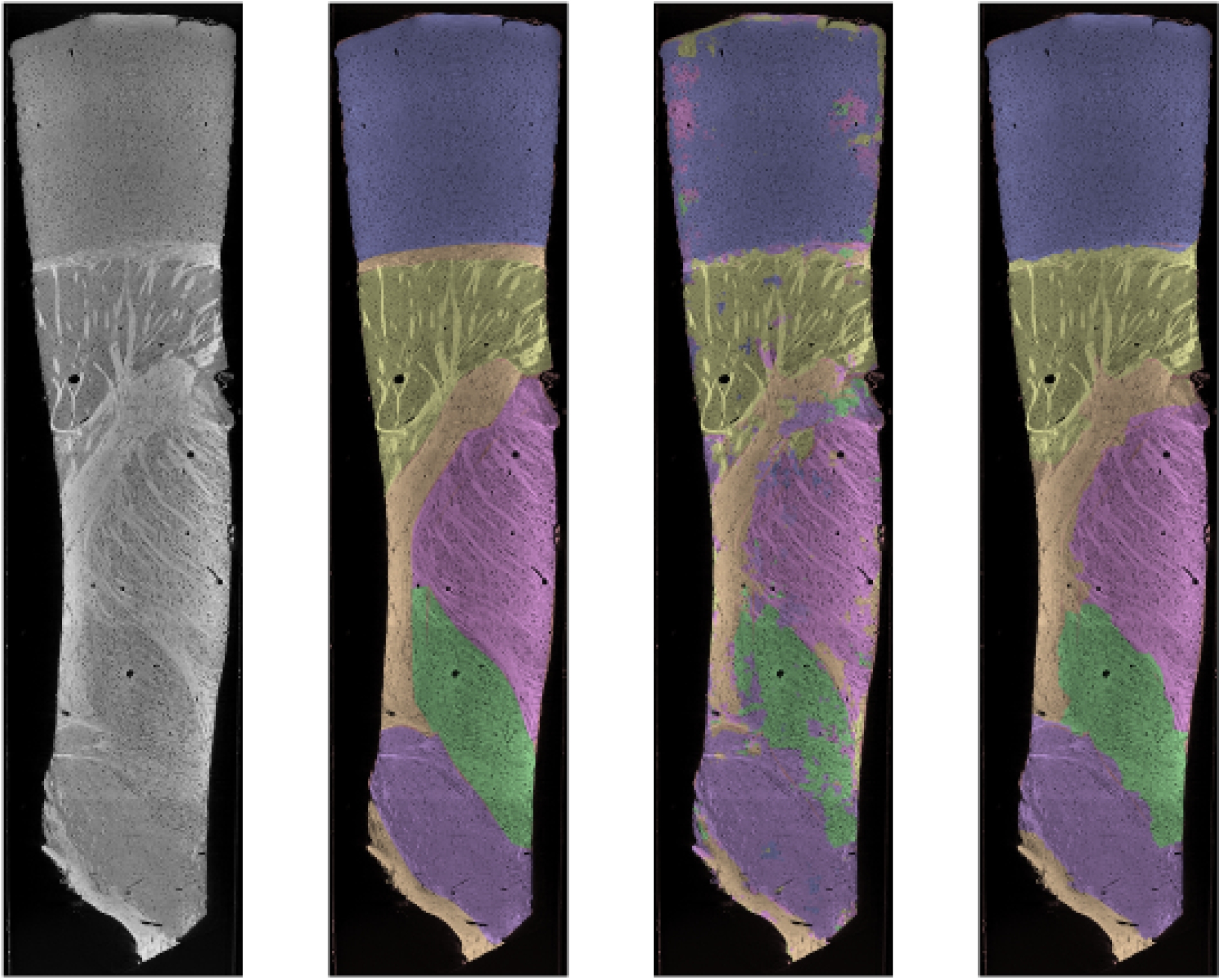
Results of using the trained CNN for large scale brain area segmentation. Shown above from left to right are: 1) Raw data of our test slice (z = 259, 100*µ*m away from the training slice), 2) Annotations for the test slice (z = 259), 3) Segmentation results obtained by passing all the extracted patches through the trained CNN (accuracy = 0.7996), and, 4) Results of applying a nearest-neighbours based clean-up algorithm (Algorithm 1) on the output of the CNN (accuracy = 0.9350).

**Figure S3:**
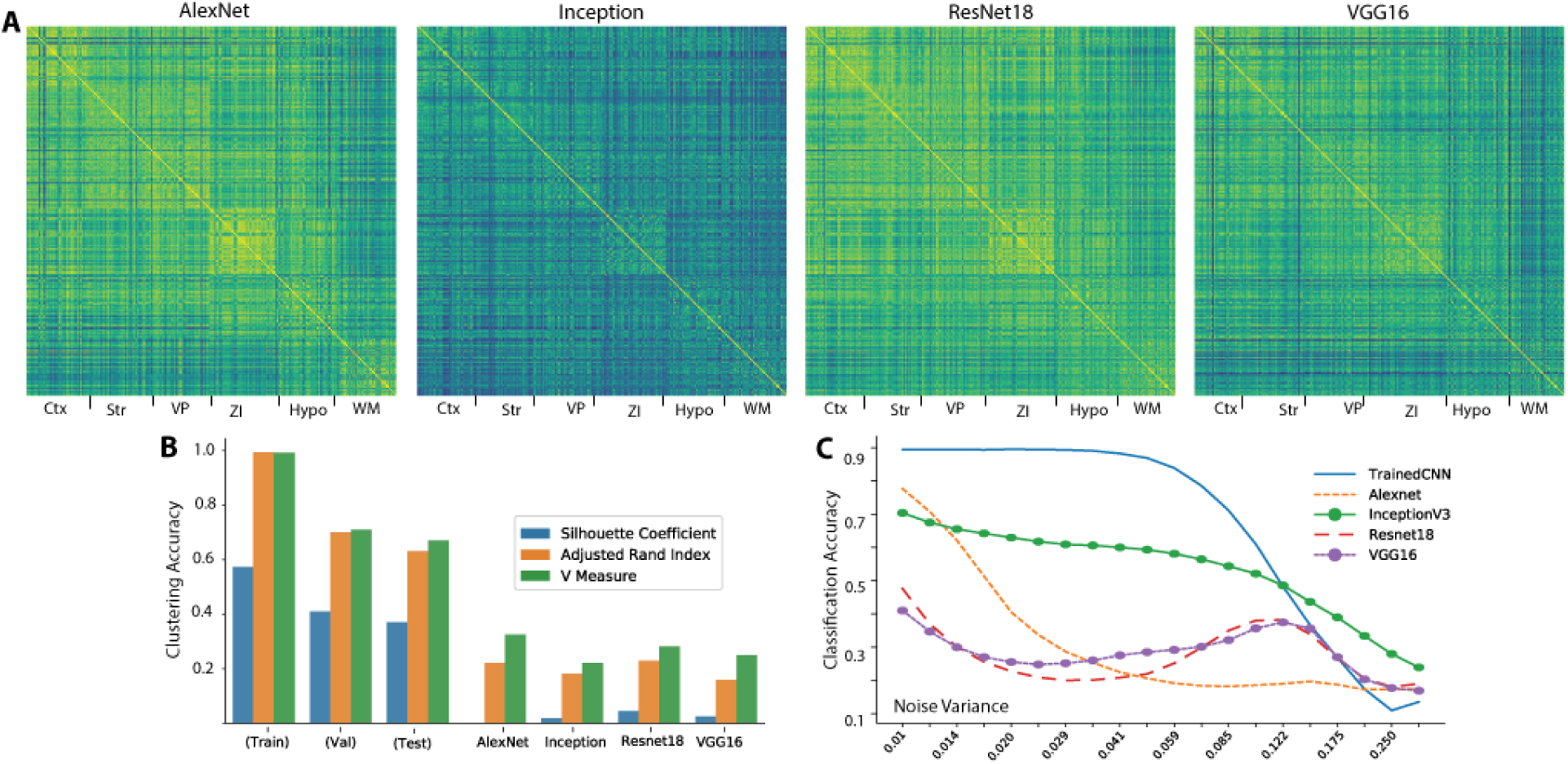
Comparison between learned features and those extracted from popular pre-trained networks. To study how discriminative features from different pre-trained networks can be for the purposes of brain area modeling, we collected the training set’s readouts from the last hidden layer of four different networks - AlexNet [52], Inception v3 [53], ResNet18 [54] and VGG16 [55], all of which were pre-trained on the ImageNet dataset [56]. Like before, we normalized these representations to be unit norm and arranged them in a matrix of the shape (*n, d*) in which they were also ordered by area. We then computed the covariance of our different feature matrices (A). We studied the quality of the representations extracted from the different networks as features for downstream tasks. To check their suitability as features for unsupervised tasks (B), we performed k-means clustering (number of clusters=6) on the extracted representations and recorded three clustering metrics, viz. the silhouette coefficient, adjusted rand index (ARI) and the v-measure. We further checked the representations’ suitability as features for a supervised brain area classification task (C) by training a logistic regression classifier on representations extracted by passing image patches from the training dataset through the different networks. We then passed patches from the test set through the networks and tested the logistic regression classifier’s ability to classify the data using the resultant features. We extended the analysis by testing the different networks’ features’ robustness to varying amounts of noise by adding zero mean Gaussian noise to the test patches before passing them through the network.

**Figure S4:**
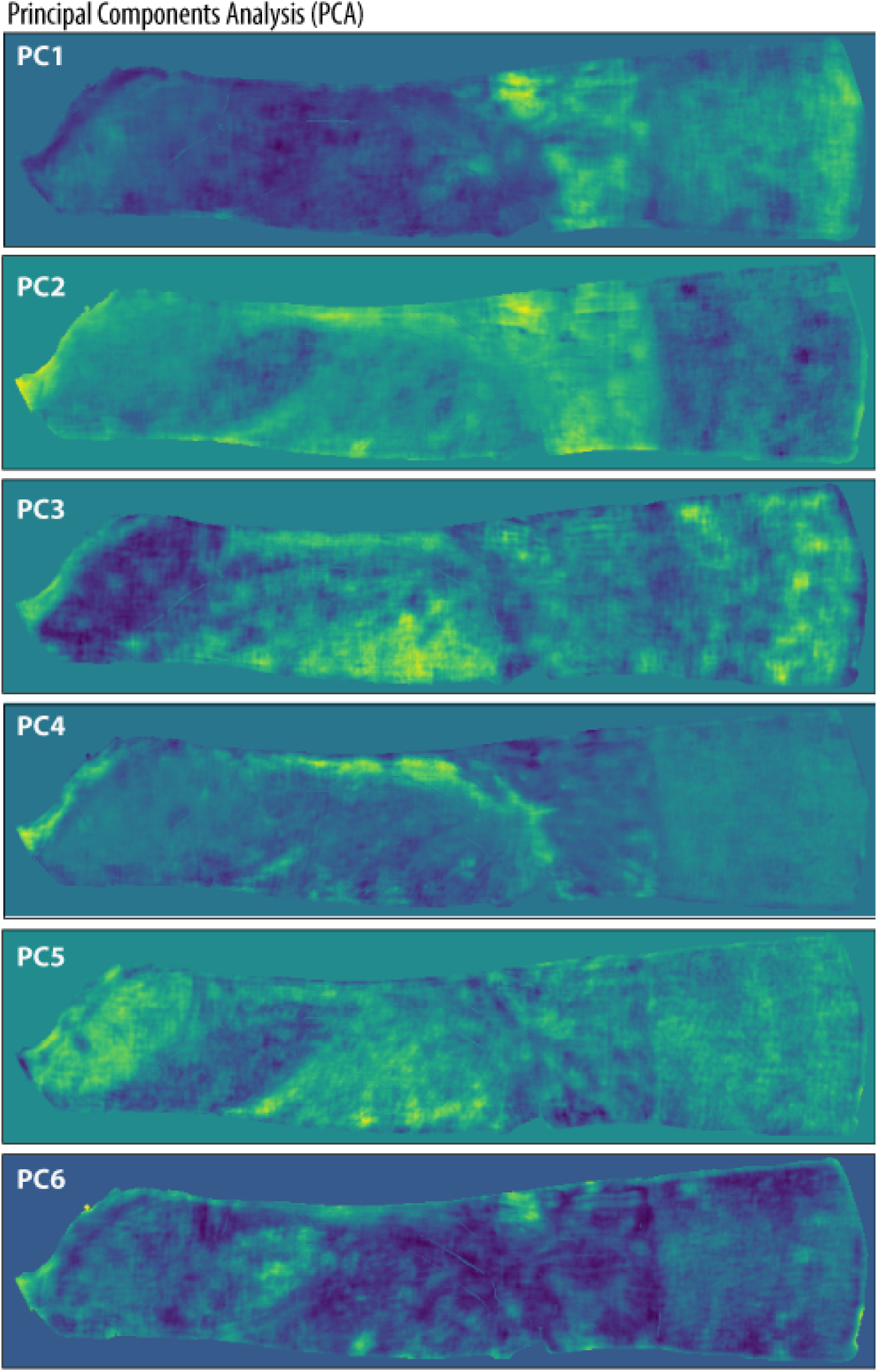
Principal components analysis applied to an entire image slice from a thalamocortical section spanning six brain areas. Embeddings of feature representations along the top six PCs. In the top two PCs: cortex and striatum, are differentiated, and then with the top three PCs, layers of cortex and VP are further differentiated. PC4 appears to strongly highlight white matter but not areas where there is a strong presence of axons (e.g., striatum). ZI appears to be distinguished with PC 5-6 - and hypothalamus and VP are also highlighted in PC5.

**Figure S5:**
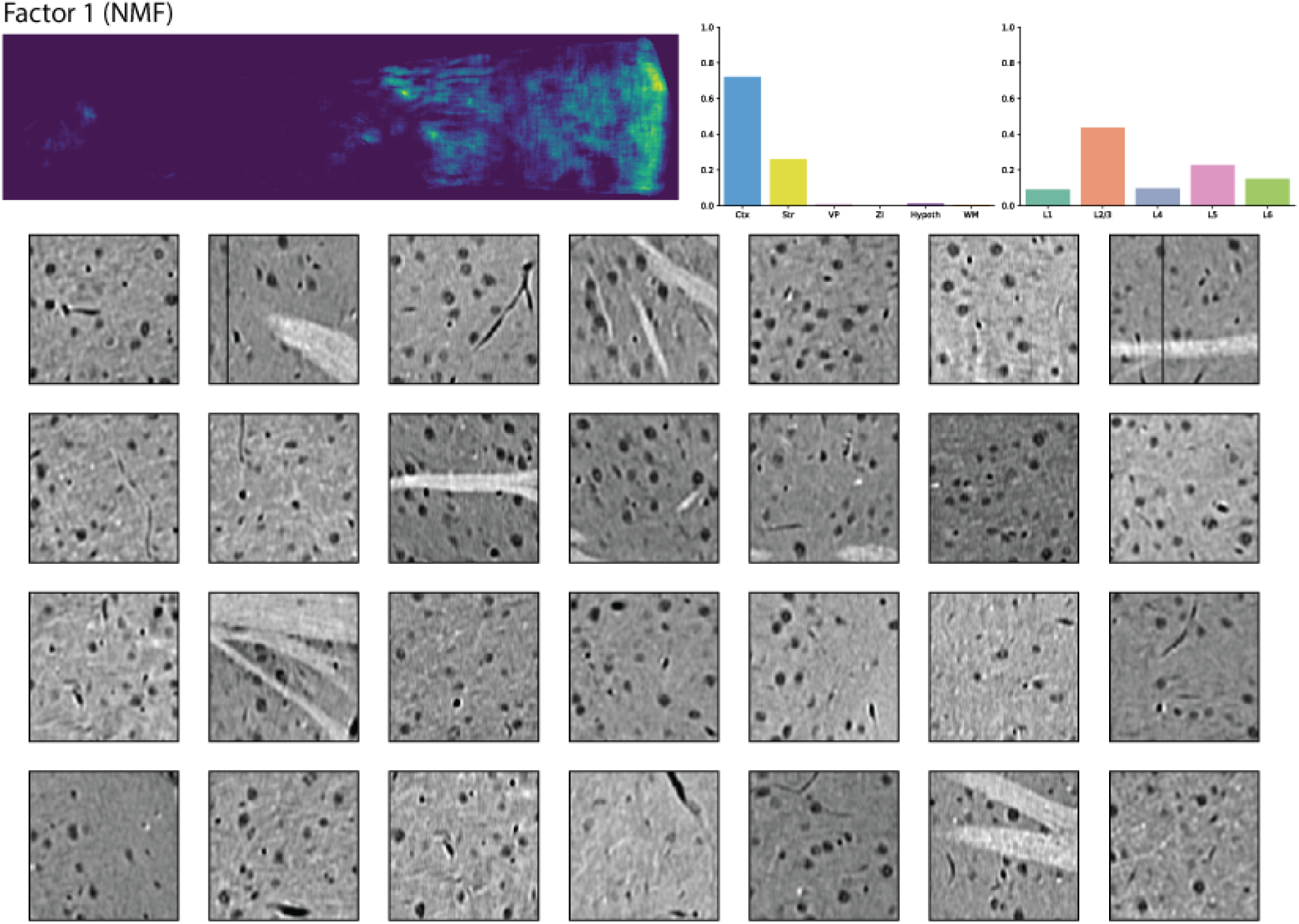
Embedding of slice in Factor 1 obtained with non-negative matrix factorization (NMF). Factor 1 strongly aligns with the Cortex and Striatum and none of the other areas. The most aligned image patches show a relatively homogeneous background, with a fair number of cells, and in the case of patches from the Striatum, some axons. The signal is spread fairly evenly in different layers of the cortex.

**Figure S6:**
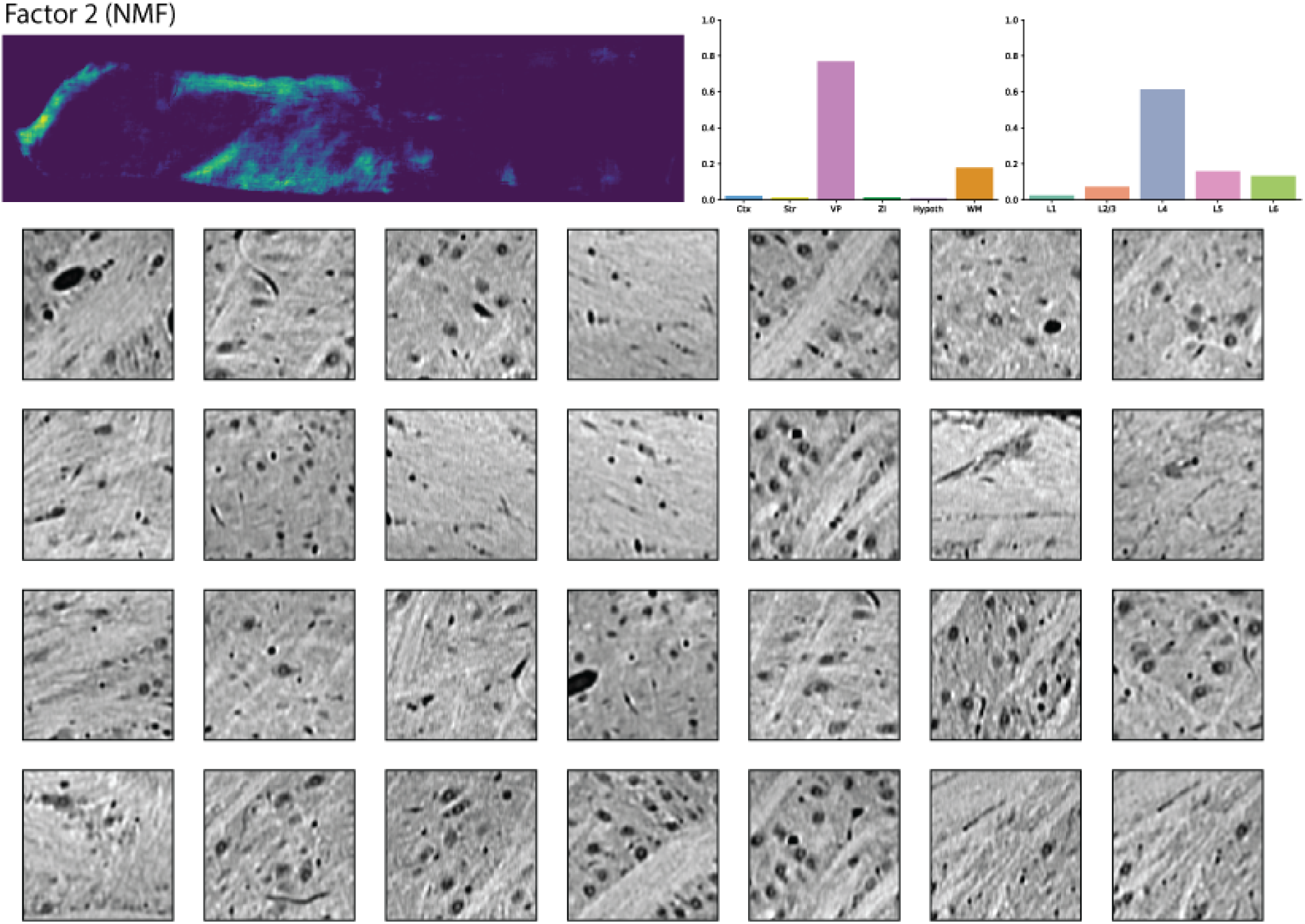
Embedding of slice in Factor 2 obtained with non-negative matrix factorization (NMF). The second non-negative factor activates axon-rich regions such as the VP and the white matter tracts. Some parts of the cortex, specifically in layer 4 are highlighted too by the factor. All of the highly activated image patches shown in the figure reveal that this factor mostly corresponds to axons, along with some cells and a few blood vessels.

**Figure S7:**
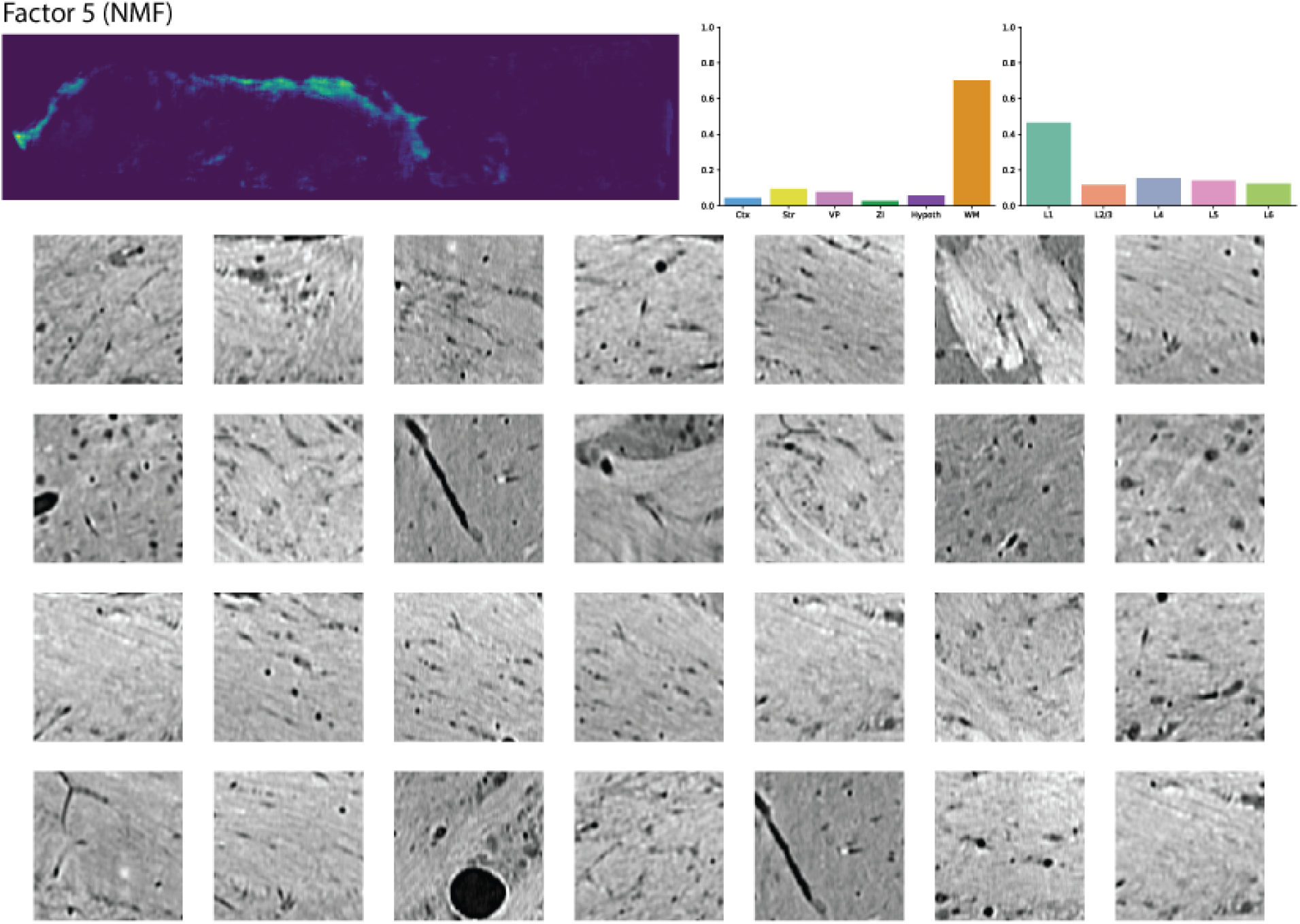
Embedding of slice in Factor 5 obtained with non-negative matrix factorization (NMF). This factor seems to align with white matter almost exclusively, and a few of the patches in VP and the Striatum are activated too. Our analysis reveals that all the top image patches that have their latents lying along this factor are predominantly made up of myelinated axons.

**Figure S8:**
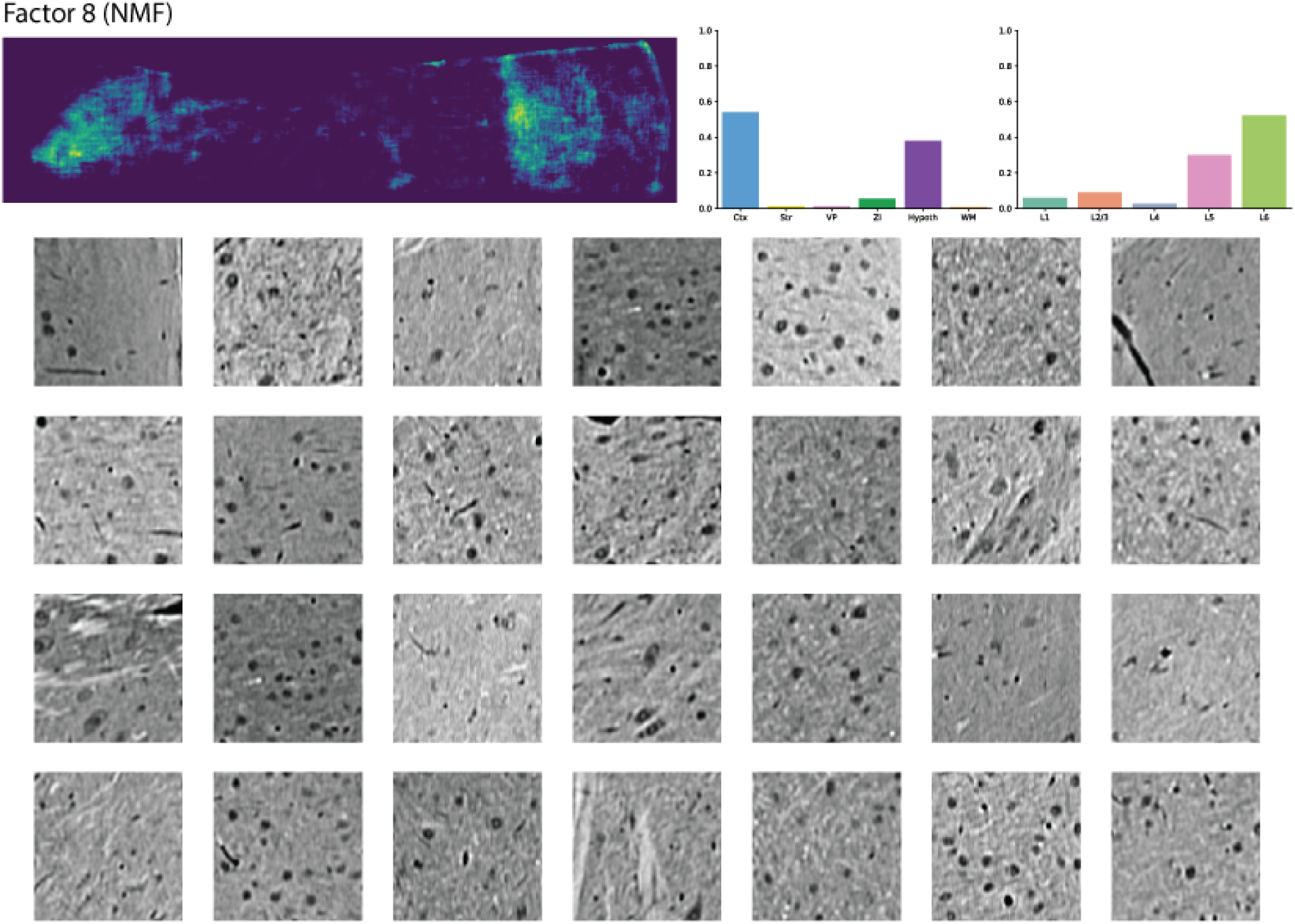
Embedding of slice in Factor 8 obtained with non-negative matrix factorization (NMF). Factor 8 aligns strongly with the cortex (especially the later layers) and hypothalamus. Examination of the top patches whose latents align with this factor reveals that these patches are mostly homogeneous, with some smattering of neural components such as cells and tiny parts of blood vessels and axons.

**Figure S9:**
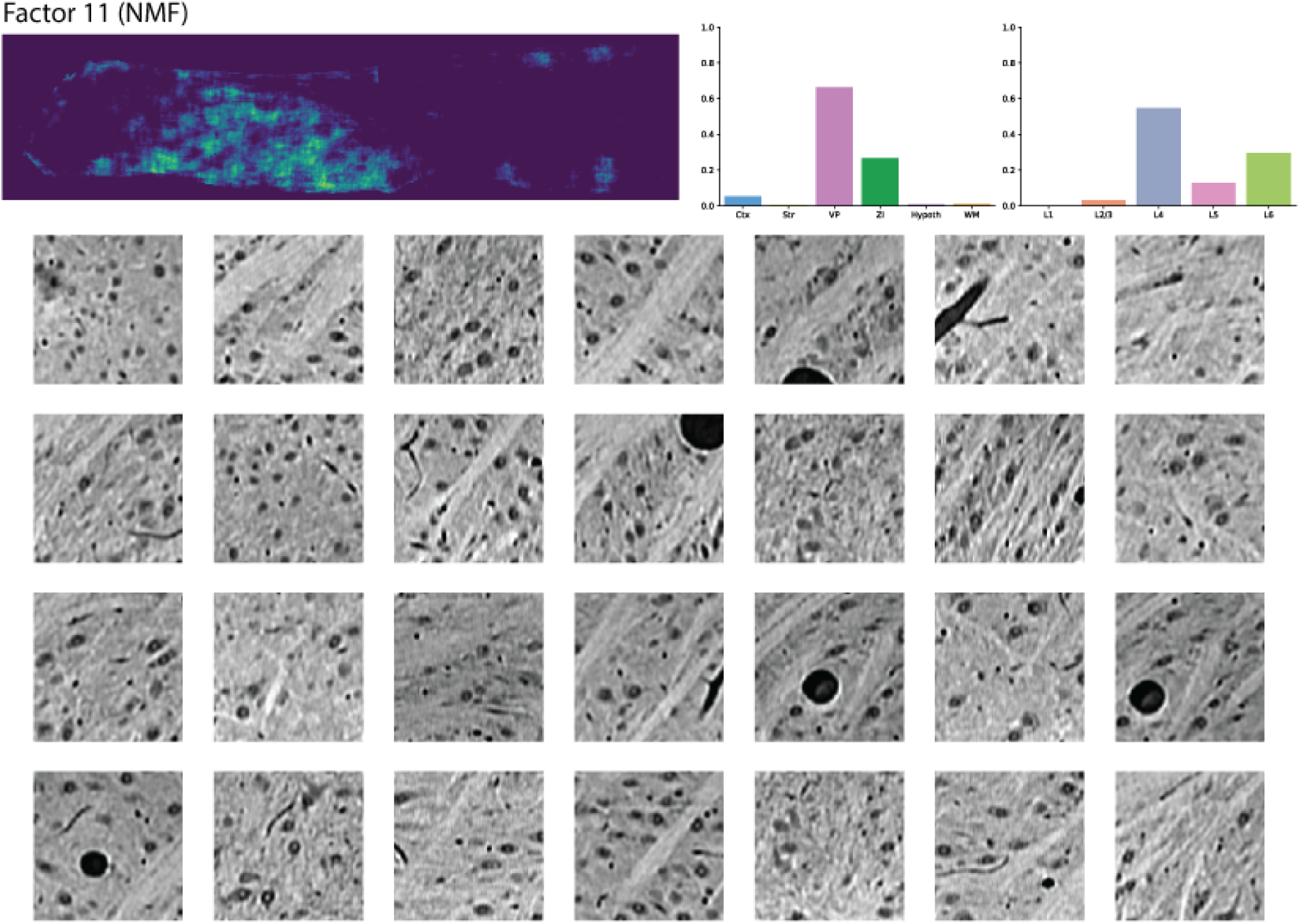
Embedding of slice in Factor 11 obtained with non-negative matrix factorization (NMF). Factor 11 highlights the VP and ZI. Some parts of the cortex align with this factor too, specifically image patches taken from regions of the cortex that seem to resemble ZI. Moreover, our analysis also reveals that the factor aligns with a fixed direction of myelinated axons.

**Figure S10:**
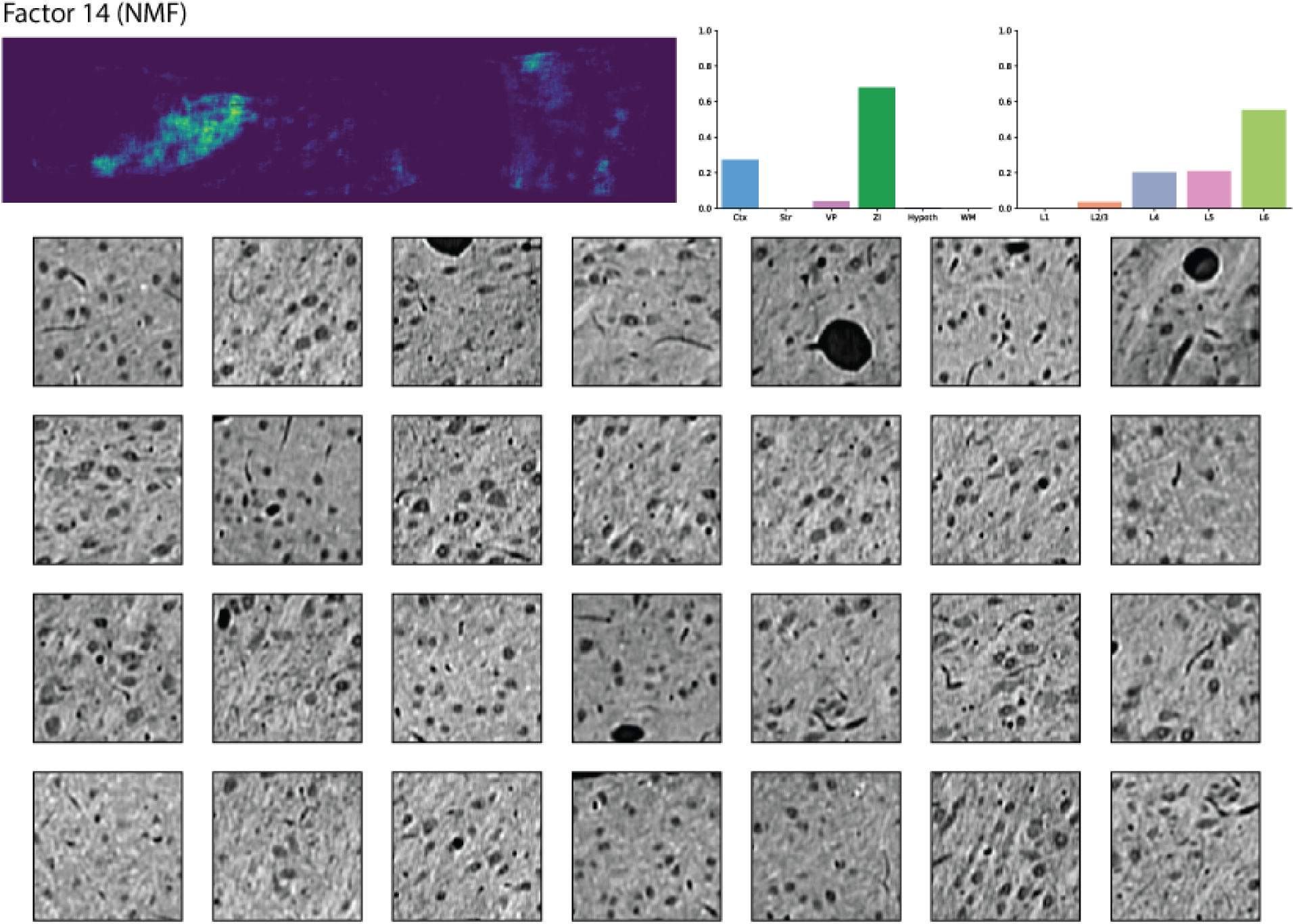
Embedding of slice in Factor 14 obtained with non-negative matrix factorization (NMF). Zona incerta is highlighted almost exclusively. This analysis also reveals that some regions of Layer 5 and 6 in cortex have some commonalities. This factor appears to reveal patterns that are unique to ZI, and also reveals some portions of cortex with diffuse axon patterns.

**Figure S11:**
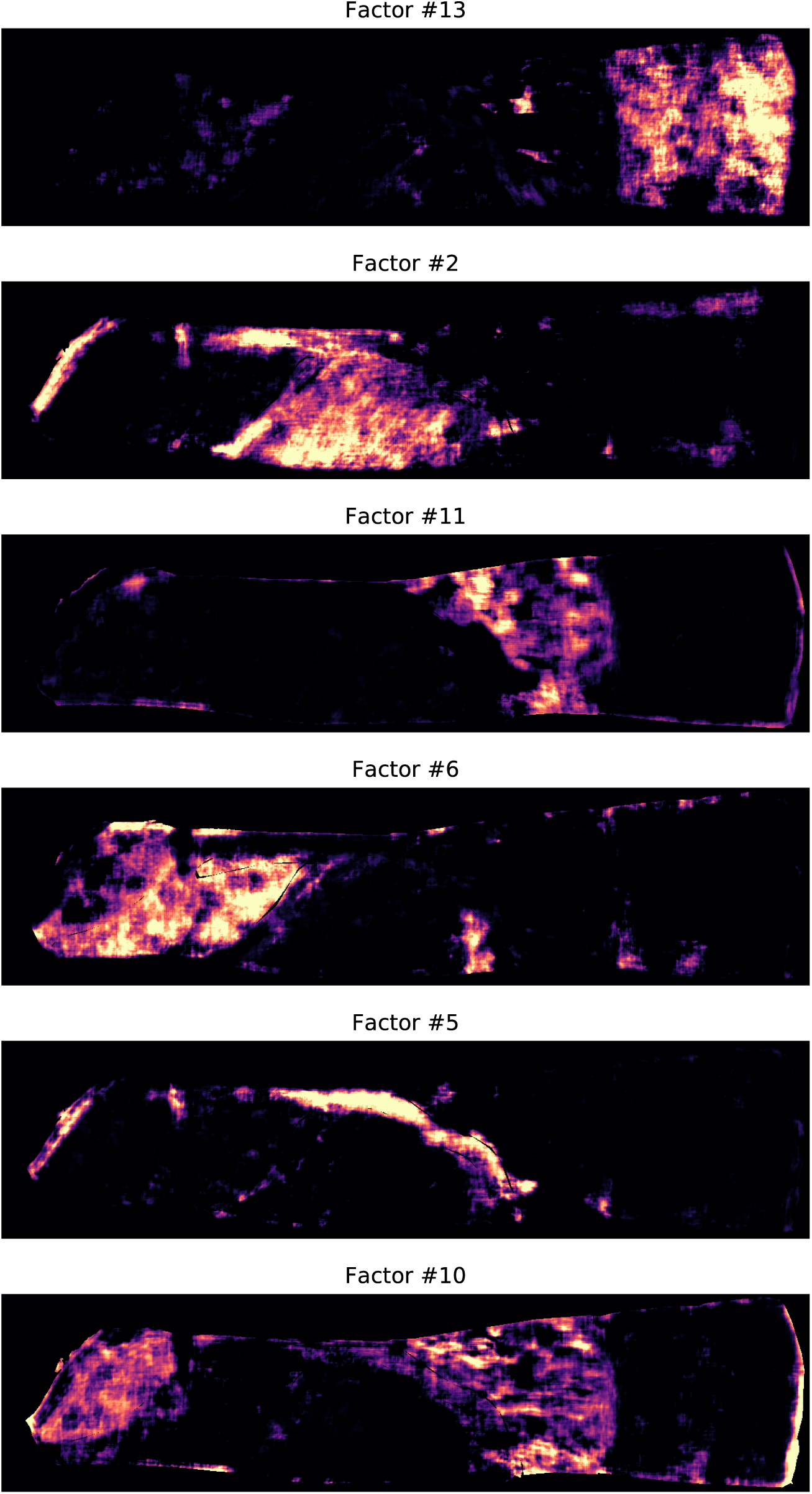
Non-negative matrix factorization results at scale for slice 0359. We show the results of NMF (*k* = 15) and our method for predictive factor sub-selection (k=6) applied to the representations collected across the entire image slice that is further away from the training slice (z = 359, ∼200*µ*m). We find that our proposed methods for microarchitecture discovery generalize fairly well, given that the NMF coefficients seem extremely similar across brain areas and exhibit almost the same inter-areal relational propoerties as those for factors of the other test slice (z = 259, Figure 4) which is ∼ 100*µ*m away.

**Figure S12:**
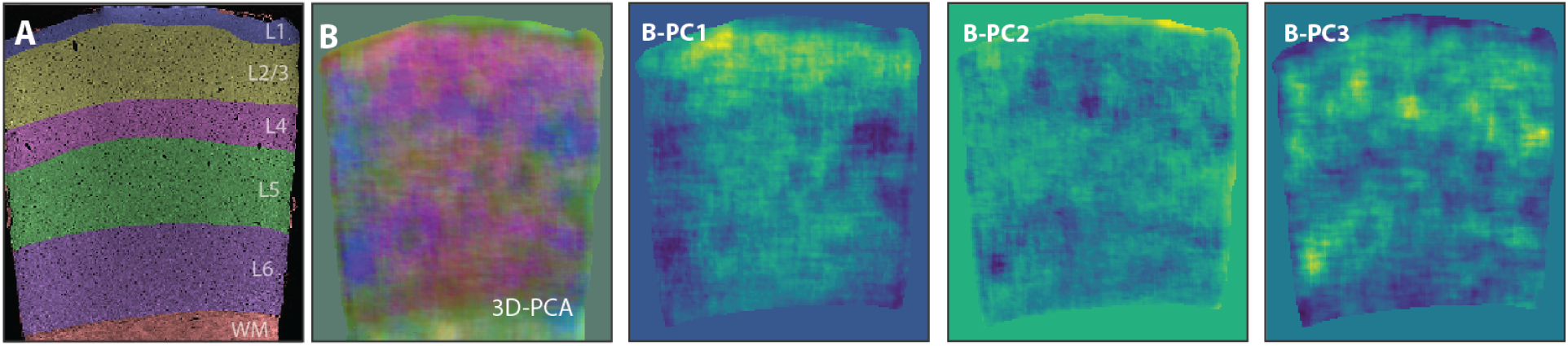
PCA embeddings of patches in cortex. To the extreme left, in (A) we provide manual annotations for layers defined in the cortex on the basis of cell density in somatosensory cortex. In (B) we show a multichannel image comprised of the coefficients of the top-3 PCs of the representations collected across the entire slice, as expressed in the cortex. The embeddings along PC1/2/3 are represented in the red/green/blue channels respectively. The three channels are displayed separately in (B-PC1/2/3).

**Figure S13:**
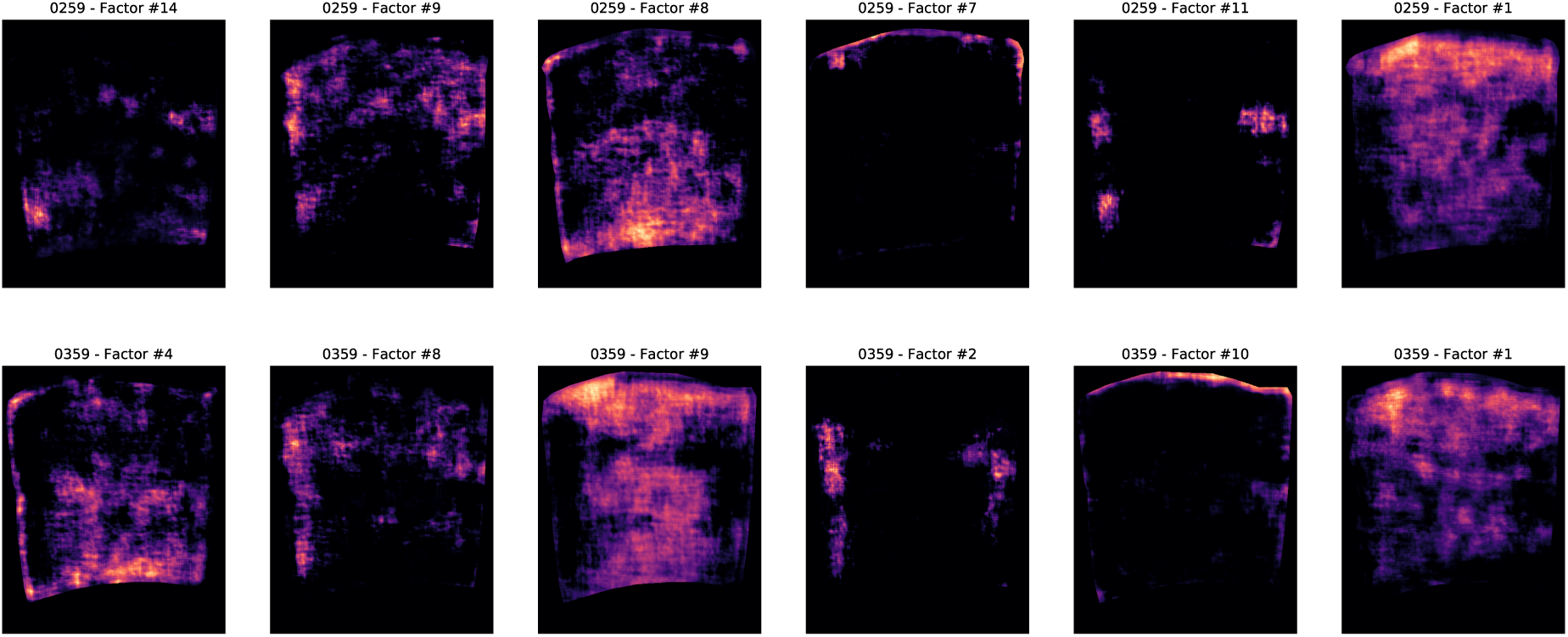
NMF coefficients of patches in cortex across multiple image slices. We compare and study the top-6 cortical factors chosen across two slices (z = 259, 359). We find that not only are the cortical expressions in across the different non-negative factors extremely similar for the two slices, but also the order in which they are chosen almost the same, speaking to the generalizability of our proposed method for brain microarchitecture discovery.

